# A hierarchical pathway for assembly of the distal appendages that organize primary cilia

**DOI:** 10.1101/2023.01.06.522944

**Authors:** Tomoharu Kanie, Julia F. Love, Saxton D. Fisher, Anna-Karin Gustavsson, Peter K. Jackson

## Abstract

Distal appendages are nine-fold symmetric blade-like structures attached to the distal end of the mother centriole. These structures are critical for formation of the primary cilium, by regulating at least four critical steps: ciliary vesicle recruitment, recruitment and initiation of intraflagellar transport (IFT), and removal of CP110. While specific proteins that localize to the distal appendages have been identified, how exactly each protein functions to achieve the multiple roles of the distal appendages is poorly understood. Here we comprehensively analyze known and newly discovered distal appendage proteins (CEP83, SCLT1, CEP164, TTBK2, FBF1, CEP89, KIZ, ANKRD26, PIDD1, LRRC45, NCS1, C3ORF14) for their precise localization, order of recruitment, and their roles in each step of cilia formation. Using CRISPR-Cas9 knockouts, we show that the order of the recruitment of the distal appendage proteins is highly interconnected and a more complex hierarchy. Our analysis highlights two protein modules, CEP83-SCLT1 and CEP164-TTBK2, as critical for structural assembly of distal appendages. Functional assay revealed that CEP89 selectively functions in RAB34^+^ ciliary vesicle recruitment, while deletion of the integral components, CEP83-SCLT1-CEP164-TTBK2, severely compromised all four steps of cilium formation. Collectively, our analyses provide a more comprehensive view of the organization and the function of the distal appendage, paving the way for molecular understanding of ciliary assembly.

## Introduction

The primary cilium is an organelle that extends from the cell surface and consists of the nine-fold microtubule-based structure (or axoneme) and the ciliary membrane (Reiter & Leroux, 2017). With specific membrane proteins (e.g., G-protein coupled receptors) accumulated on its membrane, the cilium serves as a sensor for the extracellular environmental cues (Reiter & Leroux, 2017). Biogenesis of the cilium is coupled to the cell cycle, such that the cilium mainly forms during G0/G1 phase of the cell cycle and disassembles prior to mitosis (Vorobjev & Chentsov Yu, 1982). In G0/G1 phase, the cilium extends from the mother (or older) centriole, which is distinguished from the daughter (or younger) centriole by its possession of the two centriolar substructures (Vorobjev & Chentsov Yu, 1982): distal appendages and subdistal appendages (Paintrand, Moudjou, Delacroix, & Bornens, 1992). The distal appendages are nine-fold symmetrical blade-like structures with each blade attaching to the triplet microtubules at the distal end of the mother centriole (Anderson, 1972; Bowler et al., 2019; Paintrand et al., 1992). Unlike subdistal appendages, which appear to be dispensable for the cilium formation (Mazo, Soplop, Wang, Uryu, & Tsou, 2016), the distal appendages play crucial roles in the cilium biogenesis, through their regulation of at least four different molecular steps of the cilium formation (Graser et al., 2007; Schmidt et al., 2012; Sillibourne et al., 2013; Tanos et al., 2013): 1) ciliary vesicle recruitment (Schmidt et al., 2012; Sillibourne et al., 2013), 2) recruitment of intra-flagellar transport (IFT) protein complexes (Cajanek & Nigg, 2014; Goetz, Liem, & Anderson, 2012; Schmidt et al., 2012), 3) recruitment of CEP19-RABL2 complex (Dateyama et al., 2019), which is critical for IFT initiation at the ciliary base (Kanie et al., 2017), and 4) removal of CP110 (Cajanek & Nigg, 2014; Goetz et al., 2012; Tanos et al., 2013), which is believed to suppress axonemal microtubule extension (Spektor, Tsang, Khoo, & Dynlacht, 2007), from the distal end of the mother centriole. However, how the distal appendages modulate these molecular processes is poorly understood.

To date, ten proteins were shown to localize to the distal appendages: CEP164 (Graser et al., 2007), CEP89 (also known as CCDC123) (Sillibourne et al., 2013; Sillibourne et al., 2011), CEP83 (Tanos et al., 2013), SCLT1(Tanos et al., 2013), FBF1 (Tanos et al., 2013), TTBK2 (Cajanek & Nigg, 2014), INPP5E (Q. Xu et al., 2016), LRRC45 (Kurtulmus et al., 2018), ANKRD26 (Bowler et al., 2019), and PIDD1 (Burigotto et al., 2021; Evans et al., 2021). These proteins are recruited to the distal appendages in hierarchical order, where CEP83 sits at the top of the hierarchy and recruits SCLT1 and CEP89 (Tanos et al., 2013). SCLT1 recruits CEP164 (Tanos et al., 2013), ANKRD26 (Burigotto et al., 2021; Evans et al., 2021), and LRRC45(Kurtulmus et al., 2018). CEP164 recruits TTBK2 (Cajanek & Nigg, 2014). How exactly these proteins function to organize the multiple roles of the distal appendage remains to be elucidated.

Here, we identify three more distal appendage proteins (KIZ, NCS1, and C3ORF14). The latter two will be described in an accompanying paper (Tomoharu Kanie et al., 2023). With this new set of distal appendage proteins, we sought to provide a comprehensive view of the structure, including precise localization, order of recruitment, and functional role of each distal appendage protein.

## Results

### Localization map of the new set of the distal appendage proteins

Recently, two independent studies determined the precise localization of the classical distal appendage proteins (CEP164, CEP83, SCLT1, FBF1, CEP89, TTBK2, and ANKRD26) using Stochastic Optical Reconstruction Microscopy (STORM) (Bowler et al., 2019; Yang et al., 2018). We first sought to update the localization map with the new set of distal appendage proteins using 3D-structured illumination microscopy (3D-SIM) in retinal pigment epithelial (RPE) cells. While the lateral (xy) resolution of 3D-SIM is inferior to that of STORM, the flexibility of fluorophore selection and sample preparation for multi-color imaging by 3D-SIM (Valli et al., 2021) allows us to easily locate target proteins relative to multiple centriolar markers. Using 3D-SIM, we performed three-color imaging to determine the localization of each distal appendage protein relative to CEP170, a marker for the subdistal appendage and the proximal end of the mother centriole (Sonnen, Schermelleh, Leonhardt, & Nigg, 2012), as well as the well-characterized distal appendage protein, CEP164, as references (Figure 1A; Figure 1-figure supplement 1A). Differential localization of each distal appendage protein relative to CEP164 was readily observed in either top (or axial) view or side (or lateral) view (Figure 1A; Figure 1-figure supplement 1A). As an example, the localization of FBF1, which was positioned between adjacent CEP164 structures seen in the axial view of the published STORM picture (which the authors thus identified FBF1 as a distal appendage matrix protein) (Yang et al., 2018), was also recapitulated in our SIM image (see FBF1 top view in Figure 1A; Figure 1-figure supplement 1). We also observed localization of CEP89 near the subdistal appendage in addition to its distal appendage localization (CEP89 side view in Figure 1A), consistent with the previous report (Chong et al., 2020; Yang et al., 2018). Although localization of the classical distal appendage proteins was essentially the same as that shown by two-color direct STORM imaging (Yang et al., 2018), there was one notable difference. CEP83 was shown to localize to the innermost position of the distal appendage (Bowler et al., 2019; Yang et al., 2018). We recapitulated this localization (see CEP83 inner ring in Figure 1A; Figure 1B) with the antibody used in the previous two papers (Bowler et al., 2019; Yang et al., 2018), which recognizes the C-terminal region of CEP83 (Figure 1-figure supplement 2A). The peak-to-peak diameter of the inner CEP83 ring (308.6±4.9 nm, Figure 1C) was comparable to the previous report (313±20 nm) (Yang et al., 2018). However, when we detected this protein with the antibody that detects the middle part of the protein (Figure 1-figure supplement 2A), we observed the ring located at the outermost part of the distal appendage with the diameter of 513.4±9.0 nm (see CEP83 outer ring in Figure 1A; Figure 1B; Figure 1C). Both antibodies recognized the specific band at 80-110 kDa that is lost in the CEP83 knockout cells (Figure 1-figure supplement 2B). One intriguing explanation for the difference in the ring diameter is that the protein might have an extended configuration that spans 100 nm (the difference in radius between inner and outer ring) in length. Human CEP83 protein (Q9Y592) is predicted to contain two conserved coiled-coil domains, one of which may span between 40-633 amino acids. Examination of the AlphaFold model (Jumper et al., 2021) of CEP83 shows a consistent highly extended alpha-helical structure, conserved among most species (Figure 1-figure supplement 3), supporting the model of CEP83 as a highly extended molecule. Since one alpha helix contains 3.6 residues with a distance of 0.15 nm per amino acid (Pauling, Corey, & Branson, 1951) and contour length of an amino acid is around 0.4 nm (Ainavarapu et al., 2007), 400 amino acids stretch of coiled-coil domain as well as a disordered region that consists of 40 amino acids (the maximum distance between antigens of the two CEP83 antibodies) could contribute at least 76 nm. Since the length of immunoglobulin G (IgG) is ∼8 nm (see Figure 2 of (Tan et al., 2008)), the primary and secondary antibodies at the two edges could contribute another ∼30 nm. The summed contribution from CEP83 and antibodies could readily account for the ∼100 nm difference in radius between the two rings, deriving from the extended structure of CEP83. This model, wherein CEP83 forms an extended backbone scaffolding the distal appendage, is attractive given that CEP83 is important for localization of all the other distal appendage proteins (Tanos et al., 2013) to different positions along the distal appendages (Figure 1B). Another explanation is that the two antibodies might detect different CEP83 isoforms. We think this less likely because we detect a single band that is lost in CEP83 knockouts using the two different antibodies (Figure 1-figure supplement 2B). We could also not identify different isoforms of CEP83 with similar size in the Uniprot protein database. For the recently identified distal appendage protein, PIDD1 (Burigotto et al., 2021; Evans et al., 2021), we observe a ring with similar diameter to CEP164 but that was displaced distally in the side view (see PIDD1 in Figure 1A; Figure 1B). This localization is consistent with the localization of its functional partner ANKRD26 (Burigotto et al., 2021; Evans et al., 2021) (Figure 1A; Figure1B). KIZ (or Kizuna) was localized to the similar position to ANKRD26-PIDD1. LRRC45 was located at the innermost region of the distal appendage with the smallest ring diameter (Figure 1A; Figure 1B; Figure 1C). This localization is similar to that of the inner ring of CEP83. INPP5E was reported to localize to the distal appendage in the cells grown with serum and redistribute to the cilium once cells form the organelle upon serum starvation (Q. Xu et al., 2016). This localization is also supported by the physical interaction between INPP5E and the distal appendage protein, CEP164 (Humbert et al., 2012). Indeed, we observed the INPP5E signal around the distal appendage protein CEP164 in cells grown with serum, however, we rarely observed a 9-fold ring (INPP5E in Figure 1A), typically observed with all other distal appendage proteins. Therefore, we were unable to measure the diameter of the ring formed by INPP5E. From this result, we think INPP5E is at least not a stable component of distal appendages and instead transiently localizes around the distal appendage. This localization pattern is similar to what was observed for ARL13B (see Figure 4A of (Yang et al., 2018)). Localization of the novel distal appendage proteins, NCS1 and C3ORF14, are described in an accompanying paper (Tomoharu Kanie et al., 2023), but the predicted location and their diameter are shown here for convenience (Figure 1B; Figure 1C).

**Figure 1.**
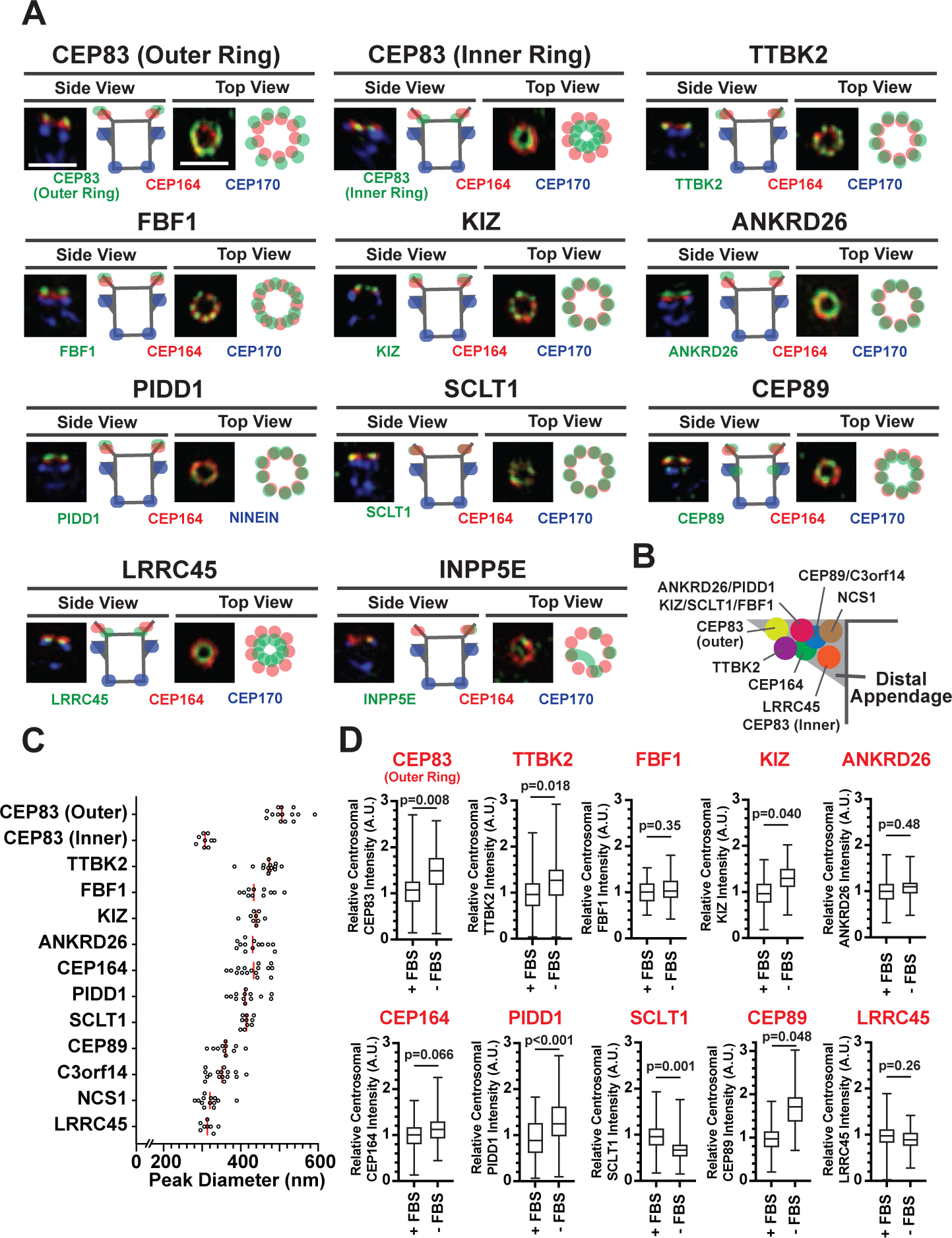
Mapping the localization of the distal appendage proteins. A. RPE cells grown to confluent in fetal bovine serum (FBS)-containing media were fixed without serum starvation (for INPP5E), or after serum starvation for 30 hours (CEP83) or 24 hours (all others). The fixed cells were stained with indicated antibodies and imaged via 3D structured illumination microscopy. Top or Side view pictures of the mother centriole are shown. The individual image is from a representative z-slice. The detailed staining and fixation condition is available in Figure1A-Source Data. Scale bar: 1 *µ*m. B. The location of each distal appendage protein on the side view of the distal appendage. The model was created from each side view shown in (A). C. The peak-to-peak diameter of each distal appendage protein. Each circle represents a measurement from each image. Red bar indicates median diameter. The raw data is available in Figure 1C-Source Data. D. Box plots showing the centrosomal signal intensity of indicated distal appendage proteins in the presence and the absence of serum. RPE cells were grown in FBS-containing media for 24 hours, and then grown in either FBS-containing media or serum free media for additional 24 hours (as shown in Figure 1-figure supplement 4A). Cells were fixed and stained with indicated antibodies. Centrosomal signal intensity of each marker was measured from fluorescent image with the method described in Materials and Methods. The relative fluorescence signal intensity compared with the average of the control is shown. A.U., arbitrary units. The data combined from three independent experiments. Statistical significance was calculated from nested T test. The raw data, sample size, experimental conditions, and detailed statistics are available in Figure 1D-Source Data.

**Figure 2.**
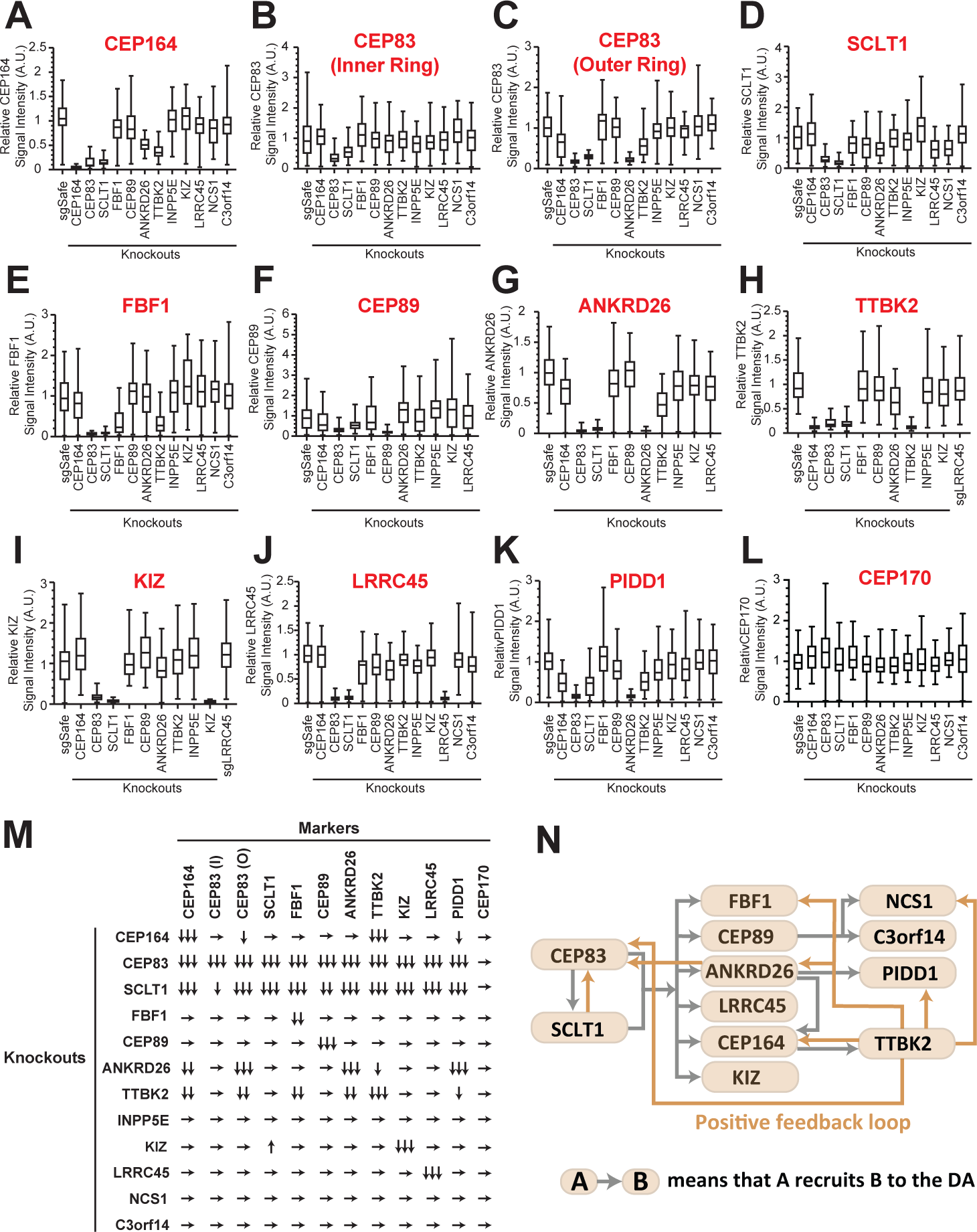
The updated hierarchy of the distal appendage proteins. A-L. Box plots showing centrosomal signal intensity of indicated distal appendage proteins (A-K) and the subdistal appendage protein, CEP170 (L) in RPE cells (control or indicated knockouts) serum-starved for 24 hours. The relative fluorescence signal intensity compared with the average of the control is shown. The data from a representative experiment. Note that FBF1 signal remains in FBF1 knockout cells, and this issue is discussed in the main text. The raw data and experimental condition are available in Figure 2A-L-Source Data. M. The summary of the signal change in each marker in indicated knockout cells compared with a control. The summary concluded from at least two independent experiments. ↓, weakly reduced; ↓↓, moderately decreased; ↓↓↓, greatly decreased or absent; ↑, weakly increased; →, unaffected. The detailed relationship between CEP89-NCS1-C3ORF14 as well as localization of each distal appendage protein in NCS1 knockout cells are available in an accompanying paper (Tomoharu Kanie, Ng, Abbott, Pongs, & Jackson, 2023). N. The updated hierarchy of the distal appendage proteins. A→B indicates that A is required for the centrosomal localization of B. CEP83 and SCLT1 is required for each other’s localization and are upstream of all the other distal appendage proteins. The outer ring, but not the inner ring, localization of CEP83 was affected in knockouts of several distal appendage proteins (ANKRD26, TTBK2, and CEP164).

Given that cilium formation of RPE cells is induced by serum deprivation (Figure 1-figure supplement 4A-B), we next tested if the localization of the distal appendage proteins changes during ciliogenesis (method described in Figure 1-figure supplement 4A; Figure 1-figure supplement 4B). Consistent with its function in ciliogenesis, we observed enhanced centriolar localization of IFT88, which requires CEP164 and TTBK2 (Goetz et al., 2012; Schmidt et al., 2012), upon serum removal (Figure 1-figure supplement 4C). Localization of one set of the distal appendage proteins (outer ring of CEP83, TTBK2, KIZ, PIDD1, and CEP89) were significantly enhanced following the serum starvation, whereas another set of distal appendage proteins (FBF1, ANKRD26, CEP164, and LRRC45) were not affected (Figure 1D). SCLT1 was the only protein that decreased its centrosomal signal upon serum deprivation.

### Updating the hierarchical map of distal appendage proteins

Distal appendage proteins are recruited to their precise location in a hierarchical order. The previous study described the order of recruitment with a simple epistatic organization, where CEP83 recruits CEP89 and SCLT1, which in turn recruits CEP164 and FBF1 (Tanos et al., 2013). With the updated set of distal appendage proteins, we sought to refine the hierarchical map of the distal appendage proteins. Notably, the original epistasis pathway using siRNA knockdowns for loss of function may fail to identify some strong requirements if limited amounts of protein are sufficient for pathway function. To this end, we generated CRISPR-Cas9 mediated knockout cells for each distal appendage protein (Figure 2-supplementary table 1A and B). We then tested for localization of each distal appendage protein in each knockout cells via immunofluorescence microscopy combined with semi-automated measurement of centrosomal signal intensity to more accurately quantify loss of localization of the proteins (see Materials and Methods). In most cases, the centrosomal signal of distal appendage proteins were barely detected in their respective knockout cells (see for example Figure 2A; Figure 2G; Figure 2I), confirming that the antibodies detect specific proteins and that the semi-automated intensity measurement was working properly. In some cases, signal was detected even in the respective knockout cells, because of the high signal observed outside of the centrosome (see for example Figure 2B), which can reflect cytoplasmic localization of the proteins or result from non-specific staining of the antibodies. FBF1 signal looked specific as it is almost completely lost in CEP83 knockout cells, however, a weak signal of FBF1was detected in FBF1 knockout cells (Figure 2E). Since both two FBF1 knockout clones had one nucleotide insertion in both alleles between coding DNA 151 and 152 (151_152insT) (Figure 2-supplementary table 1B), which results in a premature stop codon at the codon 61, we assumed that the knockout cells express truncated protein via either alternative translation or alternative splicing. We could not confirm the truncated protein because of the lack of antibody that works well for immunoblotting, and currently do not know the functional significance of the truncated proteins. Nonetheless, our semi-automated workflow provides objective and quantitative data to generate an accurate hierarchical map of the distal appendage proteins. Consistent with the previous finding (Tanos et al., 2013), the centrosomal signal intensity of all the knockout cells were greatly diminished in CEP83 knockout cells (Figure 2A-K; Figure 2M), confirming that CEP83 is the most upstream. SCLT1 depletion also showed substantial loss of localization of all the other distal appendage proteins, including CEP83 and CEP89 (Figure 2B-C; Figure 2F), suggesting that CEP83 and SCLT1 organize the distal appendage structure in a co-dependent manner. This is consistent with the previous electron micrograph showing the absence of visible distal appendages in *SCLT1*^-/-^ cells (Figure 7B of (Yang et al., 2018)). Note that SCLT1 affected the localization of the outer ring of CEP83 more strongly than that of the inner ring (Figure 2B; Figure 2C). We observed a much stronger effect of SCLT1 knockouts than the previous report (Tanos et al., 2013), which used siRNA to deplete SCLT1, likely because of the complete absence of the protein in our knockout system. Consistent with the previous report (Cajanek & Nigg, 2014), TTBK2 localization at the distal appendage was largely dependent on CEP164, as the TTBK2 signal in CEP164 knockout cells was at undetectable level similar to TTBK2 knockout cells (Figure 2H). Downstream of CEP164, TTBK2 knockout affected localization of CEP164, the outer ring of CEP83, FBF1, ANKRD26, and PIDD1 (Figure 2A; Figure 2C; Figure 2E; Figure 2G; Figure 2K; Figure 2M). The decrease of CEP164 intensity in TTBK2 knockout cells is consistent with the observation that CEP164 is a substrate of TTBK2 and that centriolar CEP164 localization was markedly increased upon overexpression of wild type but not kinase-dead TTBK2 (Cajanek & Nigg, 2014). Interestingly, localization of another TTBK2 substrate, CEP83 (Bernatik et al., 2020; Lo et al., 2019), was also affected by TTBK2 depletion (Figure 2C). Since only the outer ring of CEP83 was affected by TTBK2, we predict that CEP83 phosphorylation by TTBK2 induces conformational change of CEP83 to help enable the protein as a backbone of the distal appendage. The localization changes of FBF1, ANKRD26, and PIDD1 might suggest that these proteins may be potential substrates of TTBK2, or the localization of these proteins may be affected by phosphorylation of CEP83 or CEP164. In any case, the effect of TTBK2 depletion emphasizes that CEP164-TTBK2 complex (Cajanek & Nigg, 2014) organizes a feedback loop for other distal appendage proteins to maintain the structural integrity of the distal appendages (Figure 2N). Interestingly, ANKRD26 depletion drastically affected not only its functional partner, PIDD1 (Burigotto et al., 2021; Evans et al., 2021) (Figure 2K), but also the outer ring of CEP83 (Figure 2C). This might suggest that ANKRD26 may be crucial for maintaining the protein structure of CEP83 at the outer part of the distal appendages. The diminished CEP164 level in ANKRD26 knockout cells (Figure 2A; Figure 2M) might be explained by its direct effect or an indirect effect through the outer ring of CEP83. In contrast to a previous publication (Kurtulmus et al., 2018), we did not see the effect of LRRC45 on FBF1 localization (Figure 2E). This difference might come from the difference in the experimental setting (e.g., siRNA versus knockout). Centrosomal signal intensity of a marker of the subdistal appendage, CEP170, was not affected by any of the distal appendage proteins (Figure 2L) suggesting that distal appendage proteins are not required for the localization of subdistal appendage proteins at least in terms of signal intensity. The localization changes were mostly not due to the changes in the expression level except that protein KIZ was highly destabilized in SCLT1 knockout cells (Figure 2-figure supplement 1A).

In summary, our updated hierarchical map of distal appendage proteins shows that distal appendage proteins are highly interconnected, and the organization of the distal appendages is much more complex than what was described previously (Figure 2N). The analysis also highlighted the two modules that are critical for maintaining structural integrity of the distal appendage proteins: a CEP83-SCLT1 structural module and a CEP164-TTBK2 module providing a phosphorylation-driven positive feedback module.

### RAB34 is a superior marker for the ciliary vesicle

We next sought to understand effector functions for each distal appendage protein. The most well-established function of the distal appendages is recruitment of the ciliary vesicle (Schmidt et al., 2012), a precursor for the ciliary membrane, at the early stage of the cilium biogenesis. The recruited ciliary vesicles then fuse to form a larger vesicle through mechanisms organized by the Eps15 Homology Domain Protein 1 (EHD1) and Protein kinase C and Casein kinase 2 Substrate in Neurons (PACSIN1 and 2) proteins (Insinna et al., 2019; Lu et al., 2015). Classically, the only method to analyze ciliary vesicle recruitment to the distal appendage was elaborate electron microscopy analysis, largely because of the lack of a ciliary vesicle marker. Recently, unconventional actin-dependent motor protein, Myosin Va (MYO5A), was discovered as the earliest marker for the ciliary vesicle (C. T. Wu, Chen, & Tang, 2018). EHD1 is then recruited to the MYO5A-positive vesicle (C. T. Wu et al., 2018) to promote fusion and extension of the vesicles. However, MYO5A did not appear to be the best marker for the ciliary vesicle, because MYO5A also regulates multiple vesicle trafficking pathways including melanosome transport (X. Wu, Bowers, Rao, Wei, & Hammer, 1998) and transport of the endoplasmic reticulum (Wagner, Brenowitz, & Hammer, 2011). In agreement with the role of MYO5A outside the cilium, mutations in MYO5A cause Griscelli syndrome (Pastural et al., 1997), characterized by hypopigmentation, neurological impairment and hypotonia, characteristics distinct from other ciliopathies (Reiter & Leroux, 2017). Albinism likely reflects problems in melanosome transport. When we performed immunofluorescence microscopy, we observed a single punctum of MYO5A that colocalizes with the centrosomal marker CEP170 (Figure 3A; arrow in Figure 3-figure supplement 1A), consistent with ciliary vesicle localization (C. T. Wu et al., 2018). We also observe strong MYO5A staining surrounding the centrosome (arrowhead in Figure 3-figure supplement 1A). This pericentriolar staining persists in cells deficient in CEP83 (arrowhead in the bottom of Figure 3-figure supplement 1A), the structural component of the distal appendages (Figure 2N). Because MYO5A shows both centriolar and pericentriolar signals, it is not the best marker for ciliary vesicles.

**Figure 3.**
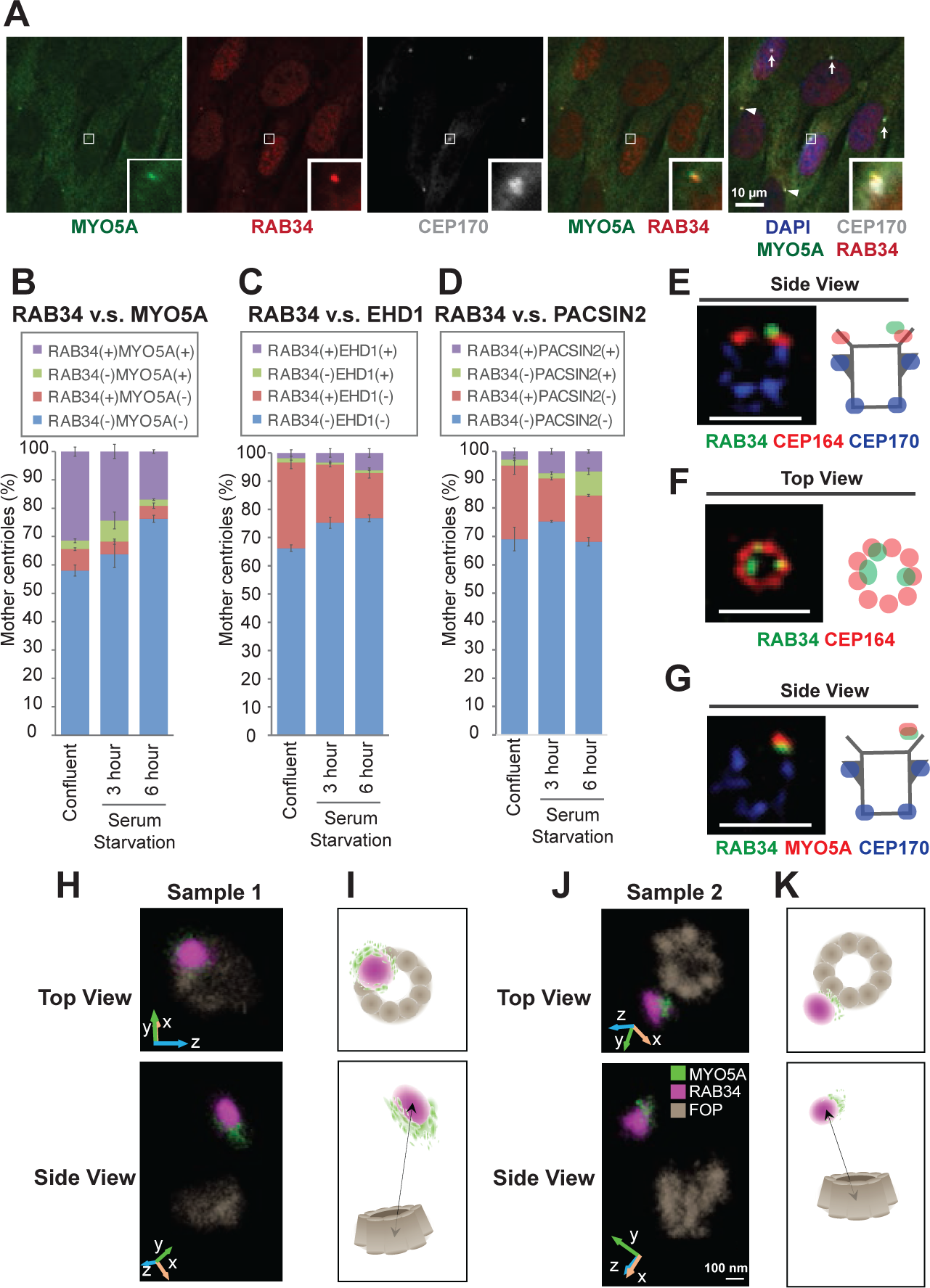
RAB34 is a marker for the ciliary vesicle. A. RPE cells were grown in 10% FBS containing media (serum-fed), fixed, stained with indicated antibodies, and imaged via wide-field microscopy. Arrows and arrowheads indicate RAB34/MYO5A negative or positive centrioles, respectively. Insets at the bottom right corner of each channel are the enlarged images of the smaller insets of each channel. Scale bar: 10 *µ*m. B-D. Quantification of the percentage of the centrioles positive for indicated markers in RPE cells grown in FBS containing media (B) or in serum-free media for 3 (C) or 6 (D) hours. Data are averaged from 3 experiments. Error bars represent ± SEM. Key statistics are available in Figure 3-figure supplement 2. The raw data, sample numbers, experimental conditions, detailed statistics are available in Figure 3B-Source data, Figure 3C-Source data and Figure 3D-Source Data. E-G. RPE cells were grown to confluent in 10% FBS containing media (serum-fed), fixed, stained with indicated antibodies, and imaged via 3D structured illumination microscopy. Scale bar: 1 *µ*m. H-K. 3D super-resolution reconstructions and illustrations of RAB34 (magenta), MYO5A (green), and FOP (gray). (H) and (J) Experimental data shown for top and side views relative to the FOP ring-structure. Orientations in the microscope 3D space are indicated by the inset axes. (I) and (K) Corresponding schematics illustrating the data and highlighting the manner in which MYO5A is located at the edge of the RAB34 distribution. FOP is here visualized with nine-fold symmetry. Arrows in the bottom panels indicate measurements of the distance of the RAB34 distribution from the mother centriole FOP structure. The schematics are not drawn to scale. Scale bar: 100 nm.

To overcome this problem, we searched for other markers for the ciliary vesicle. A recent paper suggested that the small GTPase RAB34, localizes to the ciliary vesicle (Stuck, Chong, Liao, & Pazour, 2021) and is important for ciliary vesicle recruitment/formation (S. Xu, Liu, Meng, & Wang, 2018) or for fusion of the ciliary vesicle (Ganga et al., 2021). We first tested whether RAB34 works as an early ciliary vesicle marker by staining RPE cells with RAB34, MYO5A, and a centriole marker, CEP170 (Figure 3A). We found that MYO5A and RAB34 localization are highly coupled. Most RAB34-positive centrioles have MYO5A at the centriole and vice versa (Figure 3A; Figure 3B). We confirmed the specificity of the signal using RAB34 (Figure 3-figure supplement 1B) and MYO5A knockout cells (Figure 3-figure supplement 1C). Note that the percentage of RAB34 and MYO5A double positive centrioles was highest in the confluent cells grown with serum and was decreased upon serum starvation (Figure 3B). The presence of the ciliary vesicle at the centriole before induction of cilium formation is consistent with previous electron microscopy studies (Figure 16 of (Vorobjev & Chentsov Yu, 1982) and Figure 5D of (Insinna et al., 2019)). Vesicular fusion regulators, EHD1 and PACSIN2, were recruited to the centrosome at later time points after serum withdrawal (Figure 3C; Figure 3D; Figure 3-figure supplement 2A-G), confirming that both MYO5A and RAB34 are the earliest markers of the ciliary vesicle to date. Importantly, and in contrast to MYO5A, RAB34 did not localize to the pericentriolar region (Figure 3-figure supplement 1A). This makes RAB34 a more suitable ciliary vesicle marker to assess ciliary vesicle recruitment at the distal appendages. Consistent with this, *Rab34*^-/-^ mice exhibit polydactyly and cleft palate as well as perinatal lethality, phenotypes reminiscent of cilia defects. This further supports a cilia-specific function of RAB34. In the 3D-SIM images, RAB34 localization was distal to CEP164 (Figure 3E; Figure 3-figure supplement 3A) and was displaced inwardly (Figure 3F; Figure 3-figure supplement 3F). This is reminiscent of the relative positioning between the ciliary vesicle and the distal appendages seen in electron microscopy. We also confirmed co-localization of MYO5A and RAB34 in 3D-SIM images (Figure 3G; Figure 3-figure supplement 3B-E). To more precisely define the position of RAB34 in relation to MYO5A and to the mother centriole, we turned to two-color 3D single-molecule super-resolution imaging (Bayas, Diezmann, Gustavsson, & Moerner, 2019; Bennett et al., 2020; Gustavsson, Petrov, Lee, Shechtman, & Moerner, 2018; Gustavsson, Petrov, & Moerner, 2018). This data showed how the MYO5A distribution was located at the edge of the RAB34 distribution on the vesicle (Figure 3H-K), with a 3D separation between the center of masses of the distributions of 89 nm and 67 nm in samples 1 and 2, respectively (Figure 3-figure supplement 4). We confirmed that the alternate localization of RAB34 and MYO5A is not due to channel registration by checking the complete colocalization of FOP between the two channels (Figure 3-figure supplement 5). We further confirmed this by testing colocalization of RAB34 stained with the two different secondary antibodies, Alexa Fluor 647 (AF647) and CF568 (Figure 3-figure supplement 6). The sizes of the RAB34 distributions, reflecting the measure of vesicle sizes, were found to be 230 nm x 170 nm x 190 nm for sample 1 and 190 nm x 170 nm x 250 nm for sample 2 reported as the 1/e^2^ of Gaussian fits of these distributions.

Collectively, these data suggest that RAB34 is a more specific marker for ciliary vesicle than MYO5A and is located at a distinct position from MYO5A on the ciliary vesicle.

### Distal appendages independently regulate multiple steps required for cilium formation

Distal appendages can minimally regulate four steps required for cilium formation: ciliary vesicle recruitment, IFT recruitment, IFT initiation by recruiting CEP19-RABL2, and CP110 removal. We currently do not know whether these steps are independently regulated by distal appendages or are interconnected, so that the failure of one step may interrupt the subsequent steps of the cilium formation. In the latter case, only one of the four steps may be directly regulated by the distal appendages. To test this possibility, we inhibited one step at a time and tested if other steps are affected. To inhibit ciliary vesicle recruitment, we depleted RAB34 (Figure 4-figure supplement 1A), which was shown to inhibit formation/recruitment (S. Xu et al., 2018) or fusion (Ganga et al., 2021) of the ciliary vesicle. In contrast to MYO5A depletion, which did not affect ciliogenesis, depletion of RAB34 significantly inhibited the formation of the cilium (Figure 4A), as described (Ganga et al., 2021; Oguchi, Okuyama, Homma, & Fukuda, 2020; Stuck et al., 2021; S. Xu et al., 2018).

**Figure 4.**
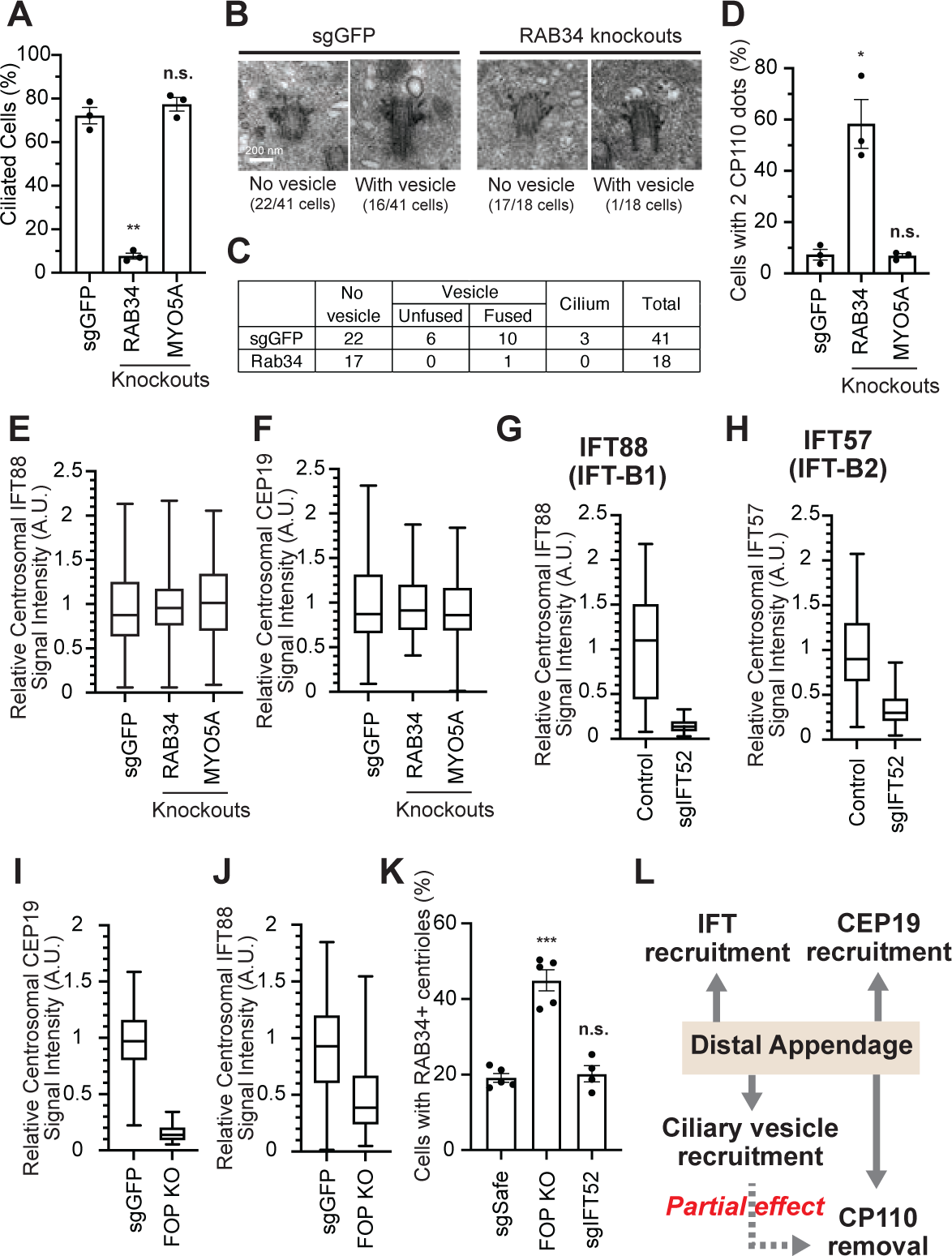
The distal appendage plays a role in ciliary vesicle recruitment, IFT recruitment, and CEP19 recruitment independently. A. Cilium formation assay in control (sgGFP), RAB34 knockout, or MYO5A knockout RPE cells serum starved for 24 hours. Data are averaged from three independent experiments, and each black dot indicates the value from an individual experiment. Error bars represent ± SEM. Statistics obtained through comparing between each knockout and control by Welch’s t-test. The raw data, experimental conditions, and detailed statistics are available in Figure 4A-Source Data. B. Transmission electron microscopy analysis of the mother centriole in control (sgGFP) or RAB34 knockout RPE cells serum starved for 3 hours. The representative images of the mother centrioles without (left) or with (right) ciliary vesicle at the distal appendages are shown. C. Quantification of the data shown in (B). The raw data and detailed statistics are available in Figure 4C-Source data. D. CP110 removal assay in control (sgGFP), RAB34 knockout, or MYO5A knockout RPE cells serum starved for 24 hours. Data are averaged from three independent experiments, and each black dot indicates the value from an individual experiment. Error bars represent ± SEM. Statistics obtained through comparing between each knockout and control by Welch’s t-test. The raw data, experimental conditions, and detailed statistics are available in Figure 4D-Source Data. E-J. Box plots showing centrosomal signal intensity of IFT88 (E, G, and J), CEP19 (F and I), or IFT57 (H) in sgGFP control (E, F, I, and J), parental RPE-BFP-Cas9 control (G and H), indicated knockouts (E, F, I, and J), or RPE cells expressing sgIFT52 (G and H) serum starved for 24 hours. At least 40 cells were analyzed per each sample. The relative fluorescence signal intensity compared with the average of the control is shown. Data from a representative experiment are shown. The raw data and experimental conditions are available in Figure 4E-J-Source Data. K. Ciliary vesicle recruitment assay in control (sgSafe) or indicated knockout RPE cells grown to confluent (without serum starvation). At least 90 cells were analyzed per each sample. The data is averaged from five independent experiments. Error bars represent ± SEM. Statistics obtained through comparing between each knockout and control by Welch’s t-test. The raw data, experimental conditions, and detailed statistics are available in Figure 4K-Source Data. L. Summary of the role of the distal appendage. The distal appendage independently regulates IFT/CEP19 recruitment and ciliary vesicle recruitment, whereas CP110 removal is partially downstream of ciliary vesicle recruitment. A.U., arbitrary units; n.s., not significant; *p < 0.05, **p < 0.01, ***p < 0.001

Electron microscopy analysis of RPE cells serum-starved for 3 hours revealed that only one out of 17 mother centrioles in RAB34-depleted cells had a ciliary vesicle, while 16 out of 41 control (sgGFP) cells had the vesicle(s) attached to centrioles (p<0.0054 in Fisher’s exact test) (Figure 4B-C). This suggests that RAB34 is important for initial recruitment/formation of the ciliary vesicle, in agreement with the previous report (S. Xu et al., 2018). Whether RAB34 is also involved in the fusion of the vesicle at the later time point after serum starvation, as shown by the other report (Ganga et al., 2021), warrant further investigation. Using RAB34 knockout cells, we tested whether disrupting the ciliary vesicle recruitment/formation affects the other steps of the cilium formation. Removal of CP110 from the mother centriole was modestly affected in RAB34 knockout cells (Figure 4D), whereas IFT and CEP19 recruitment was not affected (Figure 4E-F), suggesting that ciliary vesicle recruitment is partially important to trigger CP110 removal. This result is inconsistent with two other studies, which showed no effect on CP110 removal in RAB34 knockouts, potentially because of the difference in the duration of serum starvation (24 hours in our study versus 48 hours in other studies (Ganga et al., 2021; Stuck et al., 2021)). We next tested whether recruitment of CEP19 or IFT affects ciliary vesicle recruitment. Note that CEP19 and IFT complex proteins localize near the distal appendage in the cells grown with serum, which infrequently show primary cilia (Figure 1-figure supplement 4B), while their localization is strongly enhanced upon serum starvation (Figure 1-figure supplement 4C) (T. Kanie et al., 2017). To eliminate the IFT complexes or CEP19 from the mother centriole, we depleted either IFT52, a central component of the IFT-B complex (Taschner, Kotsis, Braeuer, Kuehn, & Lorentzen, 2014), or FGFR1OP (or FOP), which is required for centriolar localization of both IFT proteins and CEP19 (T. Kanie et al., 2017). As expected, IFT52 depletion greatly diminished the localization of all the other IFT complex proteins tested and thus inhibited the cilium formation (Figure 4G-H; Figure 4-figure supplement 2A-E). Similarly, FOP depletion abrogated the localization of CEP19 and IFT88 as well as cilium formation (Figure 4I-J, Figure 4-figure supplement 2B). Depletion of neither FOP nor IFT52 disrupted the ciliary vesicle recruitment (Figure 4K), suggesting that ciliary vesicle recruitment can proceed independently of the IFT; CEP19 pathway. We currently do not know why the number of ciliary vesicle-positive centrioles was increased in FOP knockout cells (Figure 4K). In summary, our data suggest that distal appendages independently regulate ciliary vesicle and IFT; CEP19 recruitment, whereas CP110 removal is partially downstream of ciliary vesicle recruitment (Figure 4L).

### CEP89 functions specifically in ciliary vesicle recruitment

We next sought to determine the function of each distal appendage protein. We first tested which distal appendage proteins play a role in cilium formation. The depletion of each component that is important for structural integrity of distal appendages (CEP83, SCLT1, CEP164, TTBK2) severely disrupted the cilium formation in either 24- or 48-hours serum-starved cells (Figure 5A and B). FBF1, CEP89, and ANKRD26 modestly affected cilium formation at 24 hours after serum removal (Figure 5A), but the ciliation defect in the knockout cells were ameliorated by prolonged (48 hour) serum starvation (Figure 5B). This suggests that these proteins are important for cilium formation, but that cells can compensate for the lack of these proteins to slowly catch up and form primary cilia. The distal appendage protein KIZ and LRRC45, as well as the distal appendage associated protein, INPP5E, had no effect on cilium formation (Figure 5A and B). Ciliary length was mildly affected in FBF1 and INPP5E knockout cells at the earlier time point (Figure 5-figure supplement 1A). Shorter ciliary length was observed in ANKRD26 knockout cells serum starved for either 24 or 48 hours (Figure 5-figure supplement 1A and B). Interestingly, ARL13B signal intensity inside the cilium was diminished in FBF1, CEP89, or ANKRD26 knockout cells even after the cells largely caught up on cilium formation after 48 hours of serum starvation (Figure 5-figure supplement 1C and D). This suggests that these knockouts can slowly form cilia, but the slowly formed cilia may not be functionally normal. The stronger defect in ciliary ARL13B signal in ANKRD26 might suggest a direct role of this protein in ARL13B recruitment around the distal appendages (Figure 5-figure supplement 1D).

**Figure 5.**
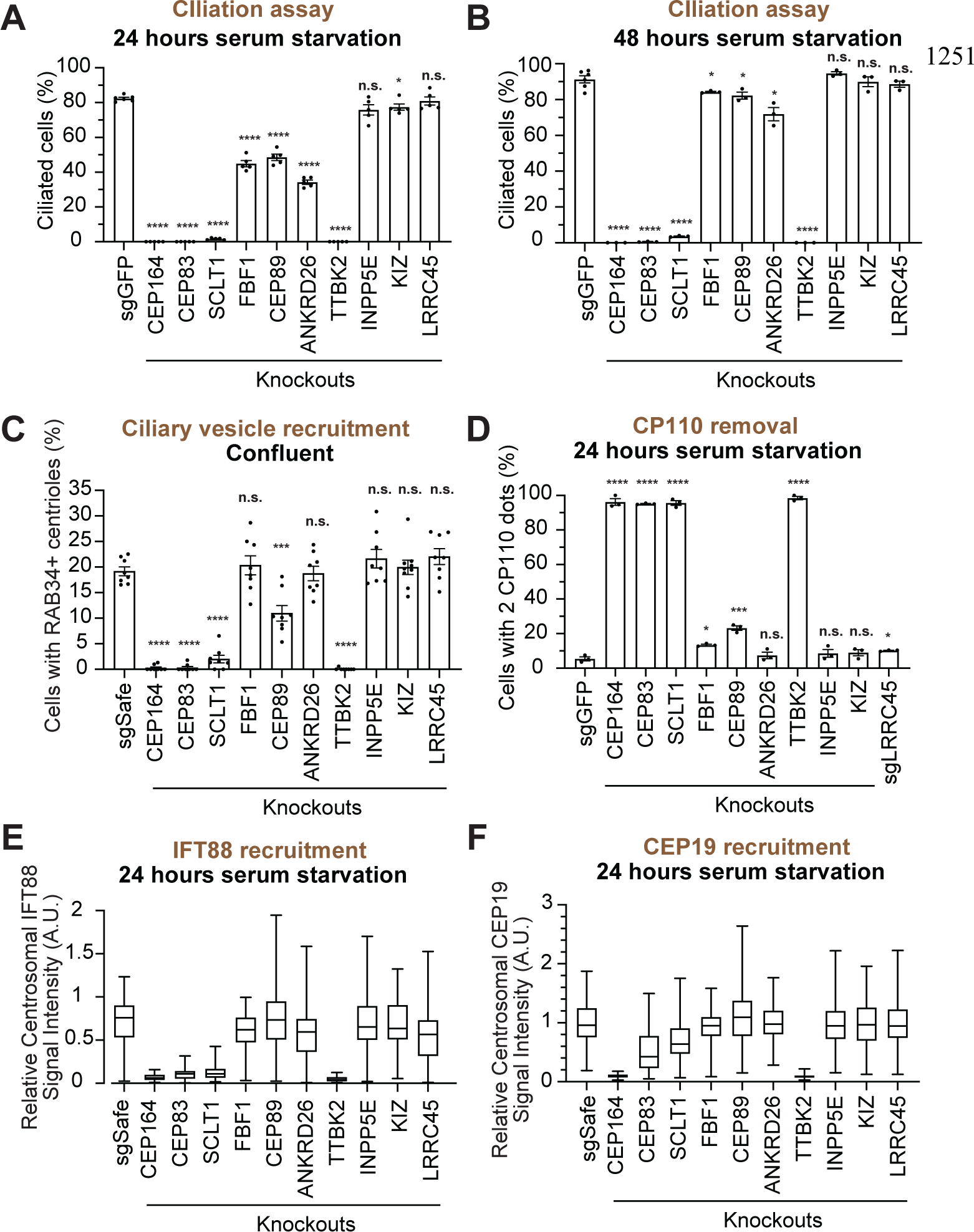
Functional analysis of distal appendage proteins reveals CEP89 as a protein important for ciliary vesicle recruitment. A-B. Cilium formation assay in control (sgGFP) and indicated knockout RPE cells serum starved for 24 hours (A) or 48 hours (B). Data are averaged from five (A) or three (B) independent experiments, and each black dot indicates the value from an individual experiment. Error bars represent ± SEM. Statistics obtained through comparing between each knockout and control by Welch’s t-test. The raw data, experimental conditions, and detailed statistics are available in Figgure 5A-B-Source Data. C. Ciliary vesicle recruitment assay in control (sgSafe) or indicated knockout RPE cells grown to confluent (without serum starvation). The data are averaged from eight independent experiments. Error bars represent ± SEM. Statistics obtained through comparing between each knockout and control by Welch’s t-test. The raw data, experimental conditions, and detailed statistics are available in Figure 5C-Source Data. D. CP110 removal assay in control (sgGFP) and indicated knockout RPE cells serum starved for 24 hours. Data are averaged from three independent experiments, and each black dot indicates the value from an individual experiment. Error bars represent ± SEM. Statistics obtained through comparing between each knockout and control by Welch’s t-test. The raw data, experimental conditions, and detailed statistics are available in Figure 5D-Source Data. E-F. Box plots showing centrosomal signal intensity of IFT88 (E) or CEP19 (F) in control (sgSafe) and indicated knockout RPE cells serum starved for 24 hours. The relative fluorescence signal intensity compared with the average of the control is shown. The data from a representative experiment are shown. The raw data and experimental conditions are available in Figure 5E-F-Source Data. A.U., arbitrary units; n.s., not significant; *p < 0.05, **p < 0.01, ***p < 0.001

We next tested the importance of each distal appendage protein in ciliary vesicle recruitment. Consistent with their critical role in the structural integrity of the distal appendages, CEP83, SCLT1, CEP164, and TTBK2 severely disrupted ciliary vesicle recruitment (Figure 5C). Interestingly, CEP89 but not the other distal appendage proteins modestly but significantly affected ciliary vesicle recruitment (Figure 5C). The importance of CEP89 in ciliary vesicle recruitment is largely consistent with a previous report (Sillibourne et al., 2013). CP110 removal was again severely affected in knockouts of the four integral component of the distal appendages (CEP83, SCLT1, CEP164, and TTBK2) (Figure 5D). CEP89 depletion partially inhibited CP110 removal (Figure 5D), correlating with the partial effect on ciliary vesicle recruitment, which is upstream of CP110 removal (Figure 4L). IFT88 recruitment was severely disturbed in the knockouts of the four integral components (CEP83, SCLT1, CEP164, and TTBK2), but not in the other knockouts (Figure 5E). The effect of CEP164-TTBK2 was slightly stronger than CEP83-SCLT1, suggesting that CEP164-TTBK2 may be more directly involved in this process.

CEP19 recruitment was strongly dependent on CEP164-TTBK2 (Figure 5F), but was only mildly affected by CEP83-SCLT1, which recruit CEP164 to the distal appendages (Figure 2A and N). This indicates that a very small amount of centriolar CEP164-TTBK2 may be sufficient to bring CEP19 near the distal appendage, and that CEP164-TTBK2, rather than CEP83-SCLT1, more directly regulates this process.

In summary, the CEP83-SCLT1 structural module brings CEP164-TTBK2 to stabilize the distal appendages, and the CEP164-TTBK2 complex plays more direct roles in the cilium formation by regulating downstream processes including ciliary vesicle recruitment, IFT; CEP19 recruitment, and CP110 removal. In contrast, CEP89 is dispensable for structural integrity of the distal appendages, but it instead plays a crucial role in the ciliary vesicle recruitment (Figure 6).

**Figure 6.**
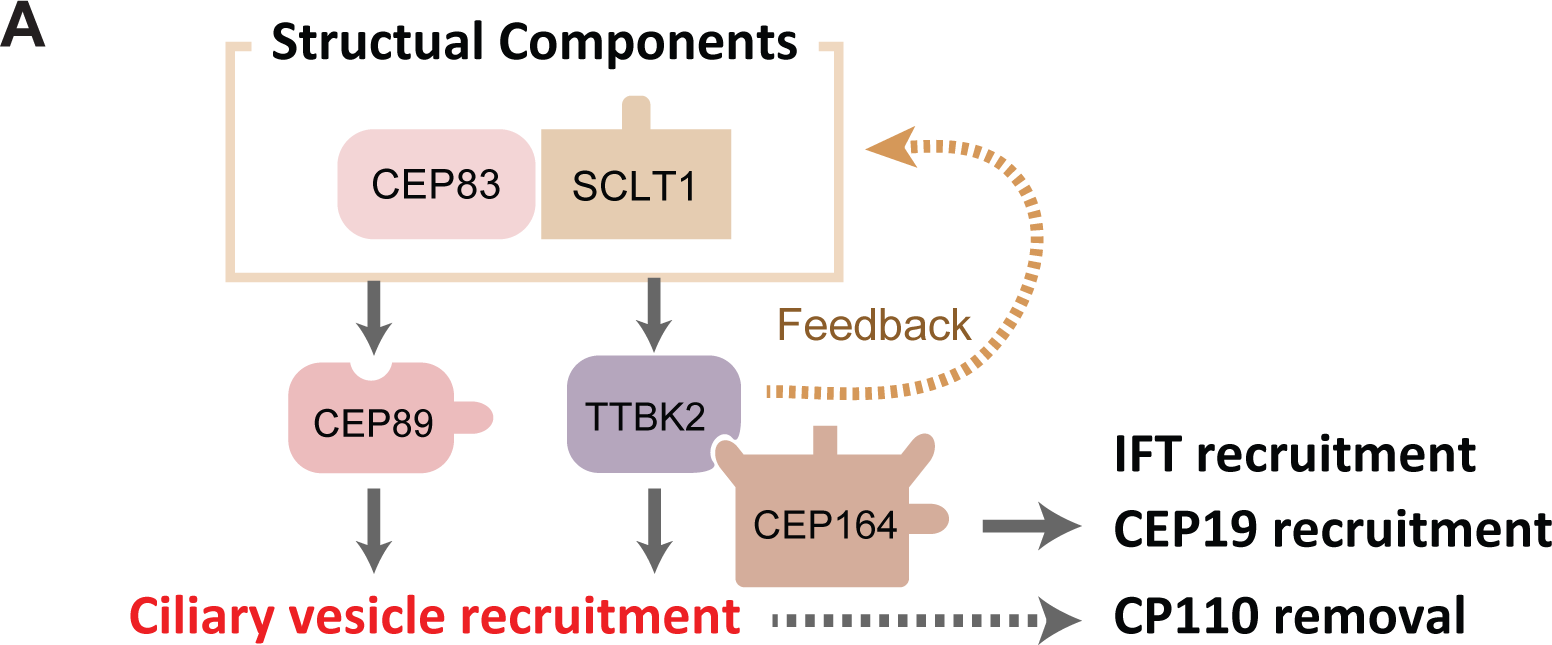
Model of the function of the distal appendage proteins. A. The CEP83-SCLT1 module serves as a structural complex, which is indispensable for the localization of all the other distal appendage proteins. TTBK2 together with its upstream protein, CEP164, is critical for proper localization of many distal appendage proteins, including the most upstream CEP83, suggesting that CEP164-TTBK2 serves as a positive feedback module. These four proteins are necessary for structural integrity of the distal appendage, thus, the lack of each protein results in severe defects in virtually all the functions of the distal appendage (IFT/CEP19/ciliary vesicle recruitment). In contrast, CEP89 is primarily important for ciliary vesicle recruitment.

## Discussion

Distal appendages are structures critical for the formation of the cilium. While their anatomical structure was described in the 1970s (Anderson, 1972; Anderson & Brenner, 1971), the first component, CEP164, was only found in 2007 (Graser et al., 2007). Since then, the list of distal appendage proteins has grown and the detailed protein architecture has been visualized mostly by super-resolution microscopy (Bowler et al., 2019; Yang et al., 2018). Nevertheless, our understanding of the function of each distal appendage protein is limited.

In this study, we sought to comprehensively characterize previously known and newly identified distal appendage proteins (KIZ, NCS1, and C3ORF14) to deepen our understanding of the distal appendages. Among the three proteins that we identified, the latter two will be described in detail in an accompanying paper (Tomoharu Kanie et al., 2023).

### The structure of the distal appendages

The distal appendages consist of twelve proteins identified so far. Each protein localizes to a different position at the distal appendages (Figure 1B). Interestingly, our current study revealed that CEP83, previously shown to locate at the innermost region of the distal appendages (Bowler et al., 2019; Yang et al., 2018), localize to the outermost region of the distal appendages when detected by antibodies that recognize a different epitope of CEP83 (Figure 1A). We currently do not know whether this is because the two antibodies detect different isoforms that differentially localize to the distal appendages or CEP83 has an extended structure that stretches the entire length of each blade of the distal appendages. The latter possibility is quite intriguing given that CEP83 is important for the localization of all the other distal appendage proteins (Figure 2) and its predicted structure, an extended alpha helix, would provide a perfect template to arrange components along the long axis of the distal appendage. Confirmation of this hypothesis needs additional investigation (e.g., crystallography and cryo-electron microscopy/tomography). SCLT1 localizes to the upper middle part of the distal appendages (Figure 1A) and was required for the localization of all the other distal appendage proteins, including the most upstream CEP83 (Figure 2M and N). This suggests that CEP83-SCLT1 module works as a structural backbone of the distal appendages. Downstream of CEP83-SCLT1, we showed that CEP164-TTBK2 complex plays an important role in maintaining the distal appendage structure. Loss of TTBK2 resulted in the decrease in the localization of several distal appendage proteins, including the most upstream CEP83 (Figure 2M and N). This is consistent with the previous finding that TTBK2 phosphorylates distal appendage proteins, such as CEP83 and CEP89 (Bernatik et al., 2020; Lo et al., 2019). Further investigations are needed to assess whether other distal appendage proteins are also phosphorylated by TTBK2 and how the phosphorylation affects the structure and function of the substrates. ANKRD26 also affected the localization of several distal appendage proteins, including the outer ring of CEP83 (Figure 2A, M, and N). These results emphasize that a set of proteins work together to organize the structure of distal appendages.

### The function of the distal appendages

Distal appendages are indispensable for cilium formation through regulation of at least four steps required for ciliogenesis: ciliary vesicle recruitment (Schmidt et al., 2012), recruitment of IFT (Schmidt et al., 2012) and CEP19-RABL2 (Dateyama et al., 2019), and CP110 removal (Goetz et al., 2012). Our functional analysis revealed that all the four steps were almost completely disrupted when each of the four critical proteins (CEP83-SCLT1-CEP164-TTBK2) was depleted. This result might simply come from disorganization of the distal appendages, or those four proteins may be directly involved in the four ciliogenic processes. Since the IFT/CEP19 recruitment defect was milder in CEP83/SCLT1 knockouts than CEP164/TTBK2 knockouts, we predict that CEP164-TTBK2 complex may play more direct roles in the IFT/CEP19 recruitment. Cajánek and Nigg created a chimeric protein that consists of the distal appendage targeting region of CEP164 and the kinase domain of TTBK2. Intriguingly, this chimera was sufficient to almost fully rescue the ciliation defect of CEP164-depleted cells (Figure 5 of (Cajanek & Nigg, 2014)), suggesting that the main function of CEP164 is recruitment of TTBK2 to the distal appendages. Since the kinase activity of TTBK2 is critical for cilium formation (Cajanek & Nigg, 2014; Goetz et al., 2012), testing whether IFT complex proteins, CEP19, or their association partners (e.g., FGFR1OP) are phosphorylation targets of TTBK2 warrant future studies.

Our study also revealed CEP89 as a protein important for ciliary vesicle recruitment, but not for other processes of cilium formation. Given that CEP89 consists of two coiled-coil domains but lacks a membrane association domain, we hypothesized that CEP89 is involved in the ciliary vesicle recruitment via its interacting partner. Indeed, our further analysis revealed that Neuronal Calcium Sensor-1 (NCS1) interacts with CEP89 and is recruited to the distal appendages by CEP89. NCS1 then captures ciliary vesicles via its myristoylation motif. This story will be described in an accompanying paper (Tomoharu Kanie et al., 2023). Importantly, CEP89 or NCS1 depletion only partially inhibits ciliary vesicle recruitment, suggesting a compensation mechanism for the recruitment. The apparent candidates that might compensate for the lack of NCS1/CEP89 are the critical distal appendage proteins, CEP83-SCLT1-CEP164-TTBK2, or their yet unknown interacting partners. This warrants future investigation.

In addition to their critical functions in cilium formation, the distal appendages seem to play important roles in other biological processes. Recent studies showed that the activation of PIDD1 requires recruitment of the protein to the distal appendages by ANKRD26. The activated PIDD1 forms a complex, called PIDDsome, with CASP2 and CRADD. The PIDDsome then cleaves MDM2 and stabilizes p53 to inhibit proliferation in response to deleterious centrosomal amplification (Burigotto et al., 2021; Evans et al., 2021). ANKRD26 apparently has a dual function: cilia-related (stabilization of the outer ring of CEP83 and controlling ciliogenesis possibly via ARL13B regulation) and cilia-independent function (PIDD1 activation). It is currently unclear why PIDD1 activation occurs at the distal appendage. Nevertheless, it is possible that other distal appendages may be involved in this process. It is also possible that distal appendages may be involved in yet unknown biological function.

Another possible role of the distal appendages is the regulation of the ciliary membrane protein composition. While there is no direct analysis, several lines of evidence support this hypothesis. First, distal appendages locate at the position where the mother centrioles attach to the plasma membrane, making it a strong candidate that modulates the composition of the ciliary membrane. Second, our current study showed that several distal appendage proteins have only modest or no effect on cilium formation (Figure 5A and B), but some of them are connected to ciliopathies (e.g., KIZ (El Shamieh et al., 2014) and ANKRD26 (Acs et al., 2015)). Finally, the previous study showed that the ciliary G-protein coupled receptor, GPR161, is detained for a short period of time in the membrane compartment likely between the transition zone and the membrane anchor point of the distal appendage (Ye, Nager, & Nachury, 2018) before going out of the cilium in response to Hedgehog activation. This implies that the distal appendages might serve as a second diffusion or trafficking barrier besides the well-established transition zone (Garcia-Gonzalo & Reiter, 2017). Future studies will test this hypothesis and define the molecular mechanisms by which the distal appendages control ciliary membrane composition.

## Supporting information

Source Data- the original files of the full raw unedited immunoblot

## Author Contributions

Conceptualization, T. K. and P. K. J.; Methodology, T. K., J. L., S. D. F., A. -K. G., and P. K. J.; Investigation, T. K., J. L., S. D. F., A. -K. G., and P. K. J.; Writing – Original Draft, T. K.; Writing – Review & Editing, T. K., J. L., S. D. F., A. -K. G., and P. K. J.; Funding Acquisition, T. K., A. -K. G., and P. K. J.; Resources, T.K., A. -K. G., and P. K. J.; Supervision, T. K., A. -K. G., and P. K. J.

## Acknowledgments

We thank Drs. Bahtiyar Kurtulmus and Gislene Pereira for LRRC45 antibodies, and Dr. Steve Caplan for the EHD1 antibody. We thank Dr. Jonathan Mulholland for technical advice on the 3D-SIM experiments. We thank Mr. John Perrino for technical support for sample preparation for the electron microscopy experiments. We thank members of the Jackson lab for helpful discussion and advice, especially Dr. Markus Kelly for establishing the method for semi-automated measurement of centrosomal signal intensity. We thank Ms. Sofía Vargas-Hernández for help developing the 3D cross-correlation code for the single-molecule data. 3D-SIM experiments were performed at the Stanford Cell Sciences Imaging Facility and were supported by Award Number 1S10OD01227601 from the National Center for Research Resources (NCRR). Electron microscopy observation was performed at the Stanford Cell Sciences Imaging Facility and were supported by NIH S10 Award Number 1S10OD028536-01, titled “OneView 4kX4k sCMOS camera for transmission electron microscopy applications”. The cell authentication service performed by MTCRO-COBRE Cell line authentication core of the University of Oklahoma Health Science Center was supported partly by P20GM103639 and National Cancer Institute Grant P30CA225520 of the National Institutes of Health (NIH). This project was supported by funds from the Baxter Laboratory for Stem Cell Research, the Stanford Department of Research, the Stanford Cancer Center, NIH grants R01GM114276 and R01GM121565 to PKJ, NIH grant P20GM103447 and a seed grant from Presbyterian Health Foundation (GRF00006006) to TK, and partial financial support from the National Institute of General Medical Sciences of the National Institutes of Health grant R00GM134187, the Welch Foundation grant C-2064-20210327, and startup funds from the Cancer Prevention and Research Institute of Texas grant RR200025 to AKG.

## Materials and Methods

### Plasmids

pMCB306, a lenti-viral vector containing loxP-mU6-sgRNAs-puro resistance-EGFP-loxP cassette, and P293 Cas9-BFP were gifts from Prof. Michael Bassik. Lenti-virus envelope and packaging vector, pCMV-VSV-G and pCMV-dR8.2 dvpr respectively, were gifts from Prof. Bob Weinberg (Addgene plasmid #8454 and #8455).

Lentiviral vectors containing single guide RNAs (sgRNAs) were generated by ligating 200 nM oligonucleotides encoding sgRNAs into the pMCB306 vector digested with BstXI (R0113S, NEB) and BlpI (R0585S, NEB) restriction enzymes. Before ligation, 4 µM of forward and reverse oligonucleotides listed in “Source Data-Primers used for genomic PCR” were annealed in 50 µl of annealing buffer (100 mM potassium acetate, 30 mM HEPES (pH7.4), and 3 mM magnesium acetate) at room temperature following denaturation in the same buffer at 95°C for 5 minutes. The targeting sequence for sgRNAs are listed in Figure 2-supplementary table 1. Gateway cloning compatible pDEST15PS vector used for bacterial protein expression was generated by inserting PreScission cleavage site immediately after GST tag into pDEST15 vector. pDEST15PS-ANKRD26 (214-537 a.a.) was generated by LR recombination between pENTR221-human ANKRD26 fragment (214-537 a.a.) and pDEST15PS vector.

### Cell line, Cell culture, Transfection, and Lentiviral expression

hTERT RPE-1 cells and 293T cells were grown in DMEM/F-12 (12400024, Thermo Fisher Scientific) supplemented with 10% FBS (100-106, Gemini), 1×GlutaMax (35050-079, Thermo Fisher Scientific), 100 U/mL Penicillin-Streptomycin (15140163, Thermo Fisher Scientific) at 37°C in 5% CO_2_. To induce cilium formation, cells were incubated in DMEM/F-12 supplemented with 1×GlutaMax and 100 U/mL Penicillin-Streptomycin (serum-free media). Both cell lines were authenticated via short-tandem-repeat based test. The authentication was performed by MTCRO-COBRE Cell line authentication core of the University of Oklahoma Health Science Center. Mycoplasma negativity of the original cell lines (hTERT RPE-1 and 293T) grown in antibiotics-free media were confirmed by a PCR based test (G238, Applied Biological Materials).

The cell lines expressing sgRNA were generated using lentivirus. Lentivirus carrying loxP-mU6-sgRNAs-puro resistance-EGFP-loxP cassette was produced by co-transfecting 293T cells with 150 ng of pCMV-VSV-G, 350 ng of pCMV-dR8.2 dvpr, and 500 ng of pMCB306 plasmids described above along with 3 µl of Fugene 6 (E2692, Promega) transfection reagent. Media was replaced 24 hr after transfection to omit transfection reagent, and virus was harvested at 48 hr post-transfection. Virus was then filtered with a 0.45 µm PVDF filter (SLHV013SL, Millipore) and mixed with 4-fold volume of fresh media containing 12.5 µg/ml polybrene (TR-1003-G, Millipore). Following infection for 66 hr, cells were selected with 10 µg/ml puromycin (P9620, SIGMA-Aldrich).

### CRISPR knockout

RPE cells expressing BFP-Cas9 were generated by infection with lentivirus carrying P293 Cas9-BFP, followed by sorting BFP-positive cells using FACSAria (BD). RPE-BFP-Cas9 cells were then infected with lentivirus carrying sgRNAs in the pMCB306 vector to generate knockout cells. After selection with 10 µg/ml puromycin, cells were subjected to immunoblotting, immunofluorescence, or genomic PCR combined with TIDE analysis (Brinkman, Chen, Amendola, & van Steensel, 2014) to determine knockout efficiency. The exact assay used for each cell line is listed in the CRISPR knockout cells summary (Figure 2-supplementary table 1). Cells were then infected with adenovirus carrying Cre-recombinase (1045N, Vector BioLabs) at a multiplicity of infection of 50 to remove the sgRNA-puromycin resistance-EGFP cassette. 10 days after adenovirus infection, GFP-negative single cells were sorted using FACSAria. The single cell clones were expanded, and their knockout efficiency were determined by immunofluorescence, immunoblot, and/or genomic PCR (the detail described in the “Figure 2-supplementary table 1”. The same number of validated single clones (typically three to four different clones) were mixed to create pooled single cell knockout clones to minimize the phenotypic variability occurred in single cell clones. The pooled clones were used in most of the experiments presented in this paper. The only exception is sgLRRC45 line used in Figure 2H, 2I, and 5D, which are RPE cells infected with sgRNA followed by removal of sgRNA-puromycin resistance-EGFP cassette and GFP-negative bulk sorting (not single cell cloning). The targeting sequences of guide RNAs are listed in the Figure 2-supplementary table 1.

### Transmission electron microscopy

Either control (sgGFP) or RAB34 knockout RPE cells were grown on 12 mm round coverslips (12-545-81, Fisher Scientific), followed by serum starvation for 3 hr. Cells were then fixed with 4% PFA (433689M, Alfa Aesar)/2% glutaraldehyde (G7526, SIGMA) in sodium cacodylate buffer (100 mM sodium cacodylate and 2 mM CaCl_2_, pH 7.4) for 1 hr at room temperature, followed by two washes with sodium cacodylate buffer. Cells were then post fixed in cold/aqueous 1% osmium tetroxide (19100, Electron Microscopy Sciences) in Milli-Q water for 1 hour at 4°C, allowed to warm to room temperature (RT) for 2 hrs rotating in a hood, and washed three times with Milli-Q water. The samples were then stained with 1% uranyl acetate in Milli-Q water at room temperature overnight. Next, the samples were dehydrated in graded ethanol (50%, 70%, 95%, and 100%), followed by infiltration in EMbed 812. Ultrathin serial sections (80 nm) were created using an UC7 (Leica, Wetzlar, Germany), and were picked up on formvar/Carbon coated 100 mesh Cu grids, stained for 40 seconds in 3.5% uranyl acetate in 50% acetone followed by staining in Sato’s Lead Citrate for 2 minutes. Electron micrographs were taken on JEOL JEM1400 (120 kV) equipped with an Orius 832 digital camera with 9 µm pixels (Gatan). To test the percentage of the ciliary vesicle positive centriole, multiple serial sections (typically 3-4) were analyzed per each mother centriole, as ciliary vesicles are often not attached to all nine blades of the distal appendage (i.e., ciliary vesicles are often not found in all the sections of the same mother centriole).

### Antibody generation

To raise rabbit polyclonal antibodies against ANKRD26, untagged human ANKRD26 fragment (214-537 a.a.) were injected into rabbits (1 mg for first injection and 500 µg for boosts). The ANKRD26 fragments were expressed as a GST fusion protein in Rosetta2 competent cells (#71402, Millipore) and purified using Glutathione Sepharose™ 4B Media (17075605, Cytiva) followed by cleavage of GST tag using GST tagged PreScission Protease (1 µg PreScission per 100 µg of recombinant protein). The ANKRD26 antibody was affinity purified from the serum with the same antigen used for injection via standard protocols.

### Immunofluorescence

For wide-field microscopy, cells were grown on acid-washed 12 mm #1.5 round coverslips (72230-10, Electron Microscopy Sciences) and fixed either in 4% paraformaldehyde (433689M, Alfa Aesar) in phosphate buffered saline (PBS) for 15 min at room temperature or in 100% methanol (A412-4, Fisher Scientific) for 5 min at -20°C. The primary antibodies used for immunofluorescence are listed in the “Source Data-List of the antibodies -Distal appendage network-”. All staining condition such as dilution of the antibodies can be found in the source data of each figure. After blocking with 5% normal serum that are matched with the species used to raise secondary antibodies (005-000-121 or 017-000-121, Jackson ImmunoResearch) in immunofluorescence (IF) buffer (3% bovine serum albumin (BP9703100, Fisher Scientific), 0.02% sodium azide (BDH7465-2, VWR International), and 0.1% NP-40 in PBS) for 30 min at room temperature, cells were incubated with primary antibody in IF buffer for at least 3 hr at room temperature, followed by rinsing with IF buffer five times. The samples were then incubated with fluorescent dye-labeled secondary antibodies (listed below) in IF buffer for 1 hr at room temperature, followed by rinsing with IF buffer five times. After nuclear staining with 4’,6-diamidino-2-phenylindole (DAPI) (40043, Biotium) in IF buffer at a final concentration of 0.5 µg/ml, coverslips were mounted with Fluoromount-G (0100-01, SouthernBiotech) onto glass slides (3050002, Epredia). Images were acquired on an Everest deconvolution workstation (Intelligent Imaging Innovations) equipped with a Zeiss Axio Imager Z1 microscope and a CoolSnap HQ cooled CCD camera (Roper Scientific). A 40x NA1.3 plan-apochromat objective lens (420762-9800, Zeiss) was used for ciliation assays, and a 63x NA1.4 plan-apochromat objective lens (420780-9900, Zeiss) was used for other analyses.

For ciliation assays, cells were plated into a 6-well plate at a density of 2 x 10^5^ cells/well and grown for 66 hr. Cells were serum starved for 24 hr unless otherwise indicated and fixed in 4% PFA. After the blocking step, cells were stained with anti-ARL13B (17711-1-AP, Proteintech), anti-CEP170 (41-3200, Invitrogen), and anti-acetylated tubulin (Ac-Tub) antibodies (T7451, SIGMA), washed, and then stained with anti-rabbit Alexa Fluor 488 (711-545-152, Jackson ImmunoResearch), goat anti-mouse IgG1-Alexa Fluor 568 (A-21124, Invitrogen), and goat anti-mouse IgG2b Alexa Fluor 647 (A-21242, Invitrogen). All the images were captured by focusing CEP170 without looking at a channel of the ciliary proteins to avoid selecting specific area based on the percentage of ciliated cells. The structures extending from the centrosome and positive for ARL13B with the length of more than 1 µm was counted as primary cilia. At least six images from different fields per sample were captured for typical analysis. Typically, at least 200 cells were analyzed per experiment. Exact number of cells that we analyzed in each sample can be found in the Source Data of corresponding figures. The percentage of ciliated cells were manually counted using the SlideBook software (Intelligent Imaging Innovations).

For ciliary vesicle assays, cells were plated into a 6-well plate at a density of 2 x 10^5^ cells/well, grown for 66 hr (without serum starvation), and fixed in 4% PFA. After the blocking step, cells were stained with anti-RAB34 (27435-1-AP, Proteintech), anti-Myosin Va (sc-365986, Santa Cruz), and anti-CEP170 (to mark centriole) antibodies (41-3200, Invitrogen), washed, then stained with goat anti-mouse IgG2a Alexa Fluor 488 (A-21131, Proteintech), goat anti-rabbit Alexa Fluor 568 (A10042, Invitrogen), and goat anti-mouse IgG1 Alexa Fluor 647 (A-21240, Invitrogen). All the images were captured by focusing CEP170 without looking at a channel of the ciliary vesicle markers to avoid selecting specific area based on the percentage of ciliary vesicle positive centrioles. At least eight images from different fields per sample were captured for typical analysis. Typically, at least 50 cells were analyzed per experiment. Exact number of cells that we analyzed in each sample can be found in the Source Data of corresponding figures.

For CP110 removal assays, cells were plated into a 6-well plate at a density of 2 x 10^5^ cells/well and grown for 66 hr. Cells were serum starved for 24 hr in 100% methanol. After the blocking step, cells were stained with anti-CP110 (12780-1-AP, Proteintech), anti-FOP (H00011116-M01, Abnova) (to mark both mother and daughter centrioles), and anti-CEP164 (sc-515403, Santa Cruz) (to mark the mother centriole) antibodies, washed, then stained with anti-rabbit Alexa Fluor 488 (711-545-152, Jackson ImmunoResearch), goat anti-mouse IgG2a-Alexa Fluor 568 (A-21134, Invitrogen), and goat anti-mouse IgG2b Alexa Fluor 647 (A-21242, Invitrogen). All the images were captured by focusing FOP without looking at a channel of the other centriolar proteins to avoid selecting specific area based on the percentage of CP110 positive centrioles. CP110 localizing to both mother and daughter centrioles (as judged by colocalization with FOP) were counted as two dots, and CP110 localizing only to daughter centriole (as judged by no colocalization with CEP164) was counted as a one dot. Exact number of cells that we analyzed in each sample can be found in the Source Data of corresponding figures.

For structured illumination microscopy, cells were grown on 18 mm square coverslips with a thickness of 0.17 mm ± 0.005 mm (474030-9000-000, Zeiss), fixed, and stained as described above. DAPI staining was not included for the structured illumination samples. Coverslips were mounted with SlowFade Gold Antifade Reagent (S36936, Life Technologies). Images were acquired on a DeltaVision OMX V4 system equipped with a 100×/1.40 NA UPLANSAPO100XO objective lens (Olympus), and 488 nm (100 mW), 561 nm (100 mW), and 642 nm (300 mW) Coherent Sapphire solid state lasers and Evolve 512 EMCCD cameras (Photometrics). Image stacks of 2 µm z-steps were taken in 0.125 µm increments to ensure Nyquist sampling. Images were then computationally reconstructed and subjected to image registration by using SoftWorx 6.5.1 software.

Secondary antibodies used for immunofluorescence were donkey anti-rabbit Alexa Fluor 488 (711-545-152, Jackson ImmunoResearch), donkey anti-mouse IgG DyLight488 (715-485-150, Jackson ImmunoResearch), goat anti-mouse IgG2a Alexa Fluor 488 (A-21131, Invitrogen), goat anti-mouse IgG1 Alexa Fluor 488 (A-21121, Invitrogen), donkey anti-rabbit IgG Alexa Fluor 568 (A10042, Invitrogen), goat anti-mouse IgG2a-Alexa Fluor 568 (A-21134, Invitrogen), goat anti-mouse IgG1-Alexa Fluor 568 (A-21124, Invitrogen), goat anti-mouse IgG2b Alexa Fluor 647 (A-21242, Invitrogen), goat anti-mouse IgG1 Alexa Fluor 647 (A-21240, Invitrogen), donkey anti-rabbit IgG Alexa Fluor 647 (711-605-152, Jackson ImmunoResearch).

### Immunolabeling and sample preparation for 3D single-molecule super-resolution imaging

For 3D single-molecule super-resolution imaging, RPE-hTERT cells were plated in the central four wells of glass-bottom chambers (µ-Slide 8 Well, Ibidi) at the density of 3×10^4^ cells/well and grown for 48 hours in DMEM/F-12 supplemented with 10% FBS, 1×GlutaMax, and 100 U/mL Penicillin-Streptomycin at 37°C in 5% CO_2_. 24 hr before fixation, the medium was replaced with fresh DMEM/F-12 supplemented with 10% FBS, 1×GlutaMax and 100 U/mL Penicillin-Streptomycin. The cells were then fixed in 100% MeOH for 5 min at -20°C. The slides were then washed twice in PBS and submerged and stored in PBS at 4°C in Samco Bio-Tite sterile containers (010002, Thermo Scientific) until the day before imaging. Cells were permeabilized with three washing steps with 0.2% (v/v) Triton-X 100 in PBS with 5 min incubation between each wash and blocked using 3% bovine serum albumin (BSA, A2058, Sigma-Aldrich) in PBS for 1 hr at room temperature. In the experiments shown in Figure 3H-K and Figure 3-figure supplement 4-5, the cells were incubated with rabbit anti-RAB34 (27435-1-AP, Proteintech, 1:500), mouse IgG2a anti-MYO5A (sc-365986, Santa Cruz, 1:1000), and mouse IgG2b anti-FOP (H00011116-M01, Abnova, 1:1000) diluted in 1% BSA in PBS at 4°C overnight, washed three times in 0.1% Triton-X 100 in PBS, and then incubated with donkey anti-rabbit Alexa Fluor 647 (ab150067, Abcam, 1:1000), goat anti-mouse IgG2b Alexa Fluor 647 (A-21242, Invitrogen, 1:1000), goat anti-mouse IgG2a CF568 (20258, Biotium, 1:1000), and goat anti-mouse IgG2b CF568 (20268, Biotium, 1:1000) diluted in 1% BSA in PBS for 1 hr shielded from light. In the experiments shown in Figure 3-figure supplement 6, the cells were incubated with rabbit anti-RAB34 (27435-1-AP, Proteintech, 1:500), mouse IgG2a anti-RAB34 (sc-365986, Santa Cruz, 1:250), and mouse IgG2b anti-FOP (H00011116-M01, Abnova, 1:1000) diluted in 1% BSA in PBS at 4°C overnight, washed three times in 0.1% Triton-X 100 in PBS, and then incubated with donkey anti-rabbit Alexa Fluor 647 (ab150067, Abcam, 1:1000), goat anti-mouse IgG2b Alexa Fluor 647 (A-21242, Invitrogen), goat anti-mouse IgG2a CF568 (20258, Biotium), and goat anti-mouse IgG2b CF568 (20268, Biotium) diluted in 1% BSA in PBS for 1 hr shielded from light. Then, the samples were washed five times with 0.1% Triton-X 100 in PBS, once with PBS and stored in PBS at 4°C while shielded from light until imaging up to several hours later. After aspiring remaining PBS, fluorescent beads (TetraSpeck, T7280, 0.2 µm, Invitrogen, diluted 1:300 in Milli-Q water) were added to each well and allowed to settle for 10 min before being washed 10x with PBS to remove unbound and loosely bound beads.

### Optical setup for 3D single-molecule super-resolution imaging

The optical setup was built around a conventional inverted microscope (IX83, Olympus) (Figure 3-figure supplement 7). Excitation lasers (560 nm and 642 nm, both 1000 mW, MPB Communications) were circularly polarized (LPVISC050-MP2 polarizers, Thorlabs; 560 nm: Z-10-A-.250-B-556 and 642 nm: Z-10-A-.250-B-647 quarter-wave plates, both Tower Optical) and filtered (560 nm: FF01-554/23-25 excitation filter, 642 nm: FF01-631/36-25 excitation filter, both Semrock), and expanded and collimated using lens telescopes. Collimated light was focused by a Köhler lens and introduced into the back port of the microscope through a Köhler lens to allow for wide-field epi-illumination. The lasers were toggled with shutters (VS14S2T1 with VMM-D3 three-channel driver, Vincent Associates Uniblitz).

The sample was positioned on an xy translation stage (M26821LOJ, Physik Instrumente) and an xyz piezoelectric stage (P-545.3C8H, Physik Instrumente). The emission from the sample was collected using a high numerical aperture (NA) objective (UPLXAPO100XO, 100x, NA 1.45, Olympus) and filtered (ZT405/488/561/640rpcV3 dichroic; ZET561NF notch filter; and ZET642NF notch filter, all Chroma) before entering a 4f imaging system. The first lens of the 4f imaging system (f = 80 mm, AC508-080-AB, Thorlabs) was placed one focal length from the intermediate image plane in the emission path. A dichroic mirror (T660lpxr-UF3, Chroma) was placed after the first 4f lens in order to split the light into two different spectral paths, where far red light (“red channel”) was transmitted into one optical path and greener light (“green channel”) was reflected into the other optical path. In order to reshape the point spread function (PSF) of the microscope to encode the axial position (z) of the individual fluorophores, transmissive dielectric double helix (DH) phase masks with ∼2 µm axial range (green channel: DH1-580-3249, red channel: DH1-680-3249, both Double Helix Optics) were placed one focal length after the first 4f lens in each path and another 4f lens was placed one focal length after the phase masks in both paths. Bandpass filters (red channel: two ET700/75m bandpass filters; green channel: ET605/70m bandpass filter, both Chroma) were placed in the paths between the phase masks and the second 4f lenses. The second 4f lenses then focused the light onto an EM-CCD camera (iXon Ultra 897, Andor) placed one focal length away from the second 4f lenses.

### Two-color 3D single-molecule super-resolution imaging

To facilitate calibration of the engineered PSFs and registration between the two channels, a solution of fiducial beads (TetraSpeck, T7280, 0.2 µm, Invitrogen) were diluted 1:5 in 10% polyvinyl alcohol (Mowiol 4-88, 17951, Polysciences Inc.) in Milli-Q water and spun-coat onto plasma-cleaned coverslips (#1.5H, 22 x 22 mm, 170 ± 5 µm, CG15CH, Thorlabs). For calibration of the PSFs, scans over a 2 µm axial range with 50 nm steps were acquired using the piezoelectric xyz translation stage. For registration measurements, the stage was translated in xy to ten different positions, and stacks of 50 frames were acquired at each position. Dark frames (400) were collected with the camera shutter closed before image acquisition, and the averaged intensity was subtracted from the calibration, registration, and single-molecule data before further analysis.

Directly prior to cell imaging, a reducing and oxygen-scavenging buffer optimized for dSTORM blinking (Halpern, Howard, & Vaughan, 2015) comprising 100 mM Tris-HCl (pH 8, J22638-K2, Thermo Scientific), 10% (w/v) glucose (215530, BD Difco), 2 µl/ml catalase (C100, Sigma-Aldrich), 560 µg/ml glucose oxidase (G2133, Sigma-Aldrich), and 143 mM β-mercaptoethanol (M6250, Sigma-Aldrich) was added to the well and the well was sealed with parafilm. The samples were then kept in this buffer both for diffraction-limited imaging and single-molecule imaging.

For diffraction-limited imaging, cells were imaged using laser intensities of 0.3 W/cm^2^ for the 642 nm laser and 1.2 W/cm^2^ for the 560 nm laser. Before beginning the single-molecule super-resolution imaging, a large fraction of the fluorophores in the field of view were converted into a dark state using 560 nm and 642 nm illumination each at ∼5 kW/cm^2^. The same laser intensities were then used for sequential acquisition of 100,000 frames of single-molecule data in each channel, first using exposure times of 50 ms for imaging of AF647 fluorophores in the red channel and then 35 ms for imaging of CF568 fluorophores in the green channel using calibrated EM gain and conversion gain of the camera of 183 and 4.41 photoelectrons / ADC count, respectively. Fiducial beads were detected in each frame to facilitate drift correction in post-processing.

### Analysis of 3D single-molecule super-resolution data

Images with acquired data from the two channels were cropped in ImageJ before analysis. Stacks that were acquired of fiducial beads for calibration were averaged over 50 frames at each unique position. These DH PSF calibration scans and the single-molecule images were used for calibration and localization using fit3Dspline in the modular analysis platform SMAP (Ries, 2020). Filter sizes and intensity count cutoffs, which serve as a threshold for template matching, were adjusted between samples based on localization previews to maximize correct localizations and minimize mislocalizations as identified by eye. Sample drift during image acquisition was accounted for by localizing fiducial beads in the same field of view as the single-molecule data. The measured motion of the fiducial bead was smoothed via cubic spline fitting (MATLAB function *csaps* with a smoothing parameter of 10^-6^) and subtracted from the single-molecule data using custom-written MATLAB scripts.

Registration between the two-color channels was completed in the x- and y-directions before correcting the z-direction. Images of dense fluorescent beads spun onto a coverslip were acquired at ten different xy positions and averaged in ImageJ. This averaged image was then cropped into the same fields of view as used for data acquisition in the two channels, and the MATLAB function *imregtform* was used to find the affine transformation that mapped the beads in the green channel onto the beads in the red channel. As the registration data was acquired at the coverslip while the single-molecule data could be acquired multiple microns above the coverslip through the cell, the registration was fine-tuned in an additional step by adapting a 2D cross-correlation approach (Schnitzbauer et al., 2018) to 3D and using it to account for any residual nanoscale offsets caused by aberrations when imaging higher up in the sample and to correct for any offset in the z-direction. The protein FOP is known to localize to the region close to the subdistal appendages of the mother centriole and daughter centriole, and forms ring-like structures (T. Kanie et al., 2017). By labeling these FOP structures with both AF647 and CF568, they served as a ground-truth for fine-tuning the channel registration. The FOP structures at the mother and daughter centrioles were manually isolated in each channel using Vutara SRX (version 7.0.00, Bruker) and cross-correlation was used to maximize the co-localization of the FOP localizations in the two registered channels (Figure 3-figure supplement 6). This cross-correlation was performed over multiple iterations until the shift between the two-color channels was below 2 nm for each axis. The translational shift applied to the red channel that yielded the maximum co-localization coefficient between the FOP structures in the two channels was then applied to all single-molecule localizations in the region of interest, thereby providing a nanoscale fine-tuning of the registration (Figure 3-figure supplement 5 and 6).

Following calibration, localization, drift correction, and registration of the data, localizations were rendered in Vutara SRX, where each localization was represented by a 3D Gaussian with 20 nm diameter and with variable opacities set to best visualize the localization density. The localizations were filtered to remove localizations with xy Cramér-Rao Lower Bound (CRLB) values from SMAP below 20 nm, and spurious localizations were removed by means of filtering for large average distance to eight nearest neighbors. This resulted in reconstructions containing the following number of localizations: sample 1, RAB34: 12276, MYO5A: 1867, and FOP: 16093; sample 2, RAB34: 7239, MYO5A: 588, and FOP: 14255. The opacities used for visualization are as follows: sample 1, RAB34: 0.03, MYO5A: 0.08, FOP: 0.05; sample 2, RAB34: 0.02, MYO5A: 0.09, FOP: 0.05 (Figure 3H and J).

Sizes of the RAB34 distributions were found by plotting histograms of the localizations along the x, y, and z axes and extracting the 1/e^2^ values from their Gaussian fits. Threshold values for the localizations to be counted as a vesicle candidate were set at 120 nm in each direction, and clusters of localizations that had smaller dimensions were excluded. This threshold was based on the dimensions of clusters of localizations many micrometers away from the centrioles that were not likely RAB34 on vesicles compared to the dimensions of vesicle candidates within a micron from the centrioles. For RAB34-MYO5A offset measurements, histograms of the localizations along the x, y, and z axes for both RAB34 and MYO5A were fitted to two Gaussians to estimate the center-of-mass (COM) separation. The difference between the centers of the two peaks yielded the offset between RAB34 and MYO5A in each dimension, and the total distance was calculated in 3D. Distances of the RAB34 and MYO5A structures from the mother centriole FOP structure were defined by the difference in 3D distance between the COM of the FOP structure and the COM of RAB34 and MYO5A, respectively.

### Immunoblot

For immunoblotting, cells were grown to confluent in a 6-well plate and lysed in 100 µl of NP-40 lysis buffer (50 mM Tris-HCl [pH7.5], 150 mM NaCl, and 0.3% NP-40 (11332473001, Roche Applied Science) containing 10 µg/ml LPC (leupeptin, pepstatin A, and chymostatin), and 1% phosphatase inhibitor cocktail 2 (P5726, SIGMA) followed by clarification of the lysate by centrifugation at 15,000 rpm (21,000 g) at 4°C for 10 min. 72.5 µl of the clarified lysates were then mixed with 25 µl of 4×Lithium Dodecyl Sulfate (LDS) buffer (424 mM Tris-HCl, 564 mM Tris-base, 8% LDS, 10% glycerol, 2.04 mM EDTA, 0.26% Brilliant Blue G250, 0.025% phenol red) and 2.5 µl of 2-mercaptoethanol (M3148, SIGMA), and incubated at 95°C for 5 min. Proteins were separated in NuPAGE™ Novex™ 4-12% Bis-Tris protein gels (WG1402BOX, Thermo Fisher Scientific) in NuPAGE™ MOPS SDS running buffer (50 mM MOPS, 50 mM Tris Base, 0.1% SDS, 1 mM EDTA, pH 7.7), then transferred onto Immobilon™-FL PVDF Transfer Membranes (IPFL00010, EMD Millipore) in Towbin Buffer (25 mM Tris, 192 mM glycine, pH 8.3). Membranes were incubated in LI-COR Odyssey Blocking Buffer (NC9232238, LI-COR) for 30 min at room temperature, and then probed overnight at 4°C with the appropriate primary antibody diluted in blocking buffer. Next, membranes were washed 3 x 5 min in TBST buffer (20 mM Tris, 150 mM NaCl, 0.1% Tween 20, pH 7.5) at room temperature, incubated with the appropriate IRDye® antibodies (LI-COR) diluted in blocking buffer for 30 min at room temperature, then washed 3 x 5 min in TBST buffer. Membranes were scanned on an Odyssey CLx Imaging System (LI-COR) and proteins were detected at wavelengths 680 and 800 nm. Primary antibodies used for immunoblotting are listed in the “Source Data-List of the antibodies -Distal appendage network-”. Secondary antibodies used for immunoblotting were IRDye® 800CW donkey anti-rabbit (926-32213, LI-COR) and IRDye® 680CW donkey anti-mouse (926-68072, LI-COR).

### Sequence alignment

Protein sequence alignment shown in Figure 1-figure supplement 2A was performed via global alignment with free end gaps using BLOSUM 62 matrix on the Geneious Prime Software.

### Experimental replicates

The term “replicates” used in this paper indicates that the same cell lines were plated at different dates for each experiment. In most cases, cell lines were thawed from liquid nitrogen at different dates and immunostaining was performed at different dates among the replicates.

### Quantification of fluorescence intensity and Statistical Analysis

#### Fluorescence intensity measurements

The fluorescence intensity was measured with 16-bit TIFF multi-color stack images acquired at 63x magnification (NA1.4) by using Image J software. All the images for measurement of centrosomal signal intensity were captured by focusing CEP170 without looking at a channel of the protein of interest (POI) to avoid selecting specific area based on the signal intensity of POI. To measure the fluorescence intensity of centrosomal proteins, channels containing CEP170 and the protein of interest (POI) were individually extracted into separate images. A rolling ball background subtraction with a rolling ball radius of 5 pixels was implemented for both CEP170 and the POI to perform local background subtraction. The mask for both CEP170 and the POI was created by setting the lower threshold to the minimum level that covers only the centrosome. Each mask was then combined by converting the two masks to a stack followed by z projection. The combo mask was then dilated until the two masks were merged. After eroding the dilated masks several times, the fluorescent intensity of the POI was measured via “analyze particles” command with optimal size and circularity. The size and circularity are optimized for individual POI to detect most of the centrosome in the image without capturing non-centrosomal structure. Outliers (likely non-centrosomal structure) were then excluded from the data using the ROUT method with a false discovery rate of 1% using GraphPad Prism 9 software to further remove signals from non-centrosomal structures, as the signals from those are typically extremely lower or higher than those from centrioles. Fluorescence intensity of ciliary proteins were measured similarly to centrosomal proteins but with several modifications. A mask was created for only ARL13B by setting the lower threshold to the minimum level that covers only cilia. The size and circularity are optimized for individual POI to detect only cilia without capturing non-ciliary structure. Image macros used for the automated measurement described above are found in the supplementary files.

To test whether the difference in the signal intensity is statistically different between control and test samples, the intensity measured through the described method was compared between control and test samples using nested one-way ANOVA with Dunnett’s multiple comparisons test if there are more than two replicates. In case there are less than three replicates, the statistical test was not performed in a single experiment, as the signal intensity is affected slightly by staining procedure and statistical significance is affected largely by the number of cells examined. For example, we saw statistical significance in the signal intensity with the same samples that are stained independently if we analyze large number of the cells (more than 100 cells). Instead, we confirmed the same tendency in the change of fluorescence intensity in the test samples across two replicates.

#### Statistical analysis for ciliation, ciliary vesicle recruitment, and CP110 removal assay

For ciliation, ciliary vesicle recruitment, and CP110 removal assay, the number of ciliated cells from the indicated number of replicates were compared between control (sgGFP or sgSafe) and the test samples using Welch’s t test. The exact number of samples and replicates are indicated in the resource data of the corresponding figures.

#### Statistical analysis for ciliary length measurements

For ciliary length measurements, shown in figure 5-figure supplement 5A and B, the ciliary length was compared between control and the knockouts using nested t test.

#### Diameter measurements of the distal appendage rings

For diameter measurements of the ring shown in Figure 1C, maximal intensity projection with top view images of the centriolar ring were first carried out and the peak-to-peak diameter was measured from four different angles and averaged to reduce the variability caused by tilting of the centriole. The number of the top view of the distal appendage rings analyzed is indicated in Figure1C-source data.

For all the statistics used in this paper, asterisks denote *: 0.01 ≤ p < 0.05, **: p < 0.01, ***: p < 0.001, n.s.: not significant. All the statistical significance was calculated by using GraphPad Prism 9 software.

#### Materials Availability Statement

All the newly created materials used in this paper including ANKRD26 antibody, plasmids, stable cell lines are readily available from the corresponding authors upon request.

## Figure supplements

**Figure 1-figure supplement 1.**
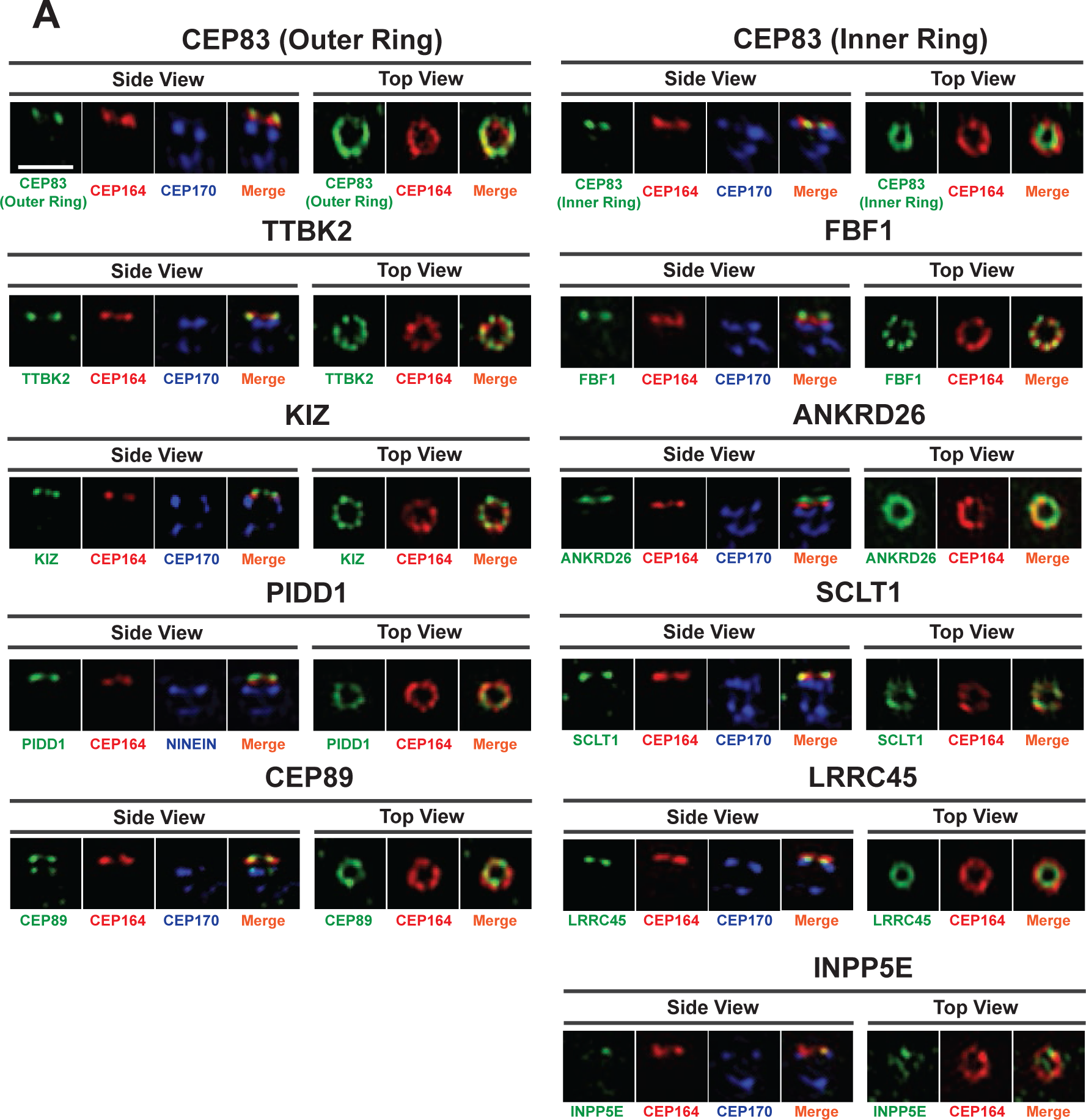
Individual channels of the images shown in Figure 1A. A. 3D structured illumination images of indicated distal appendage proteins shown in Figure 1A shown in individual channels. Scale bar: 1 *µ*m.

**Figure 1-figure supplement 2.**
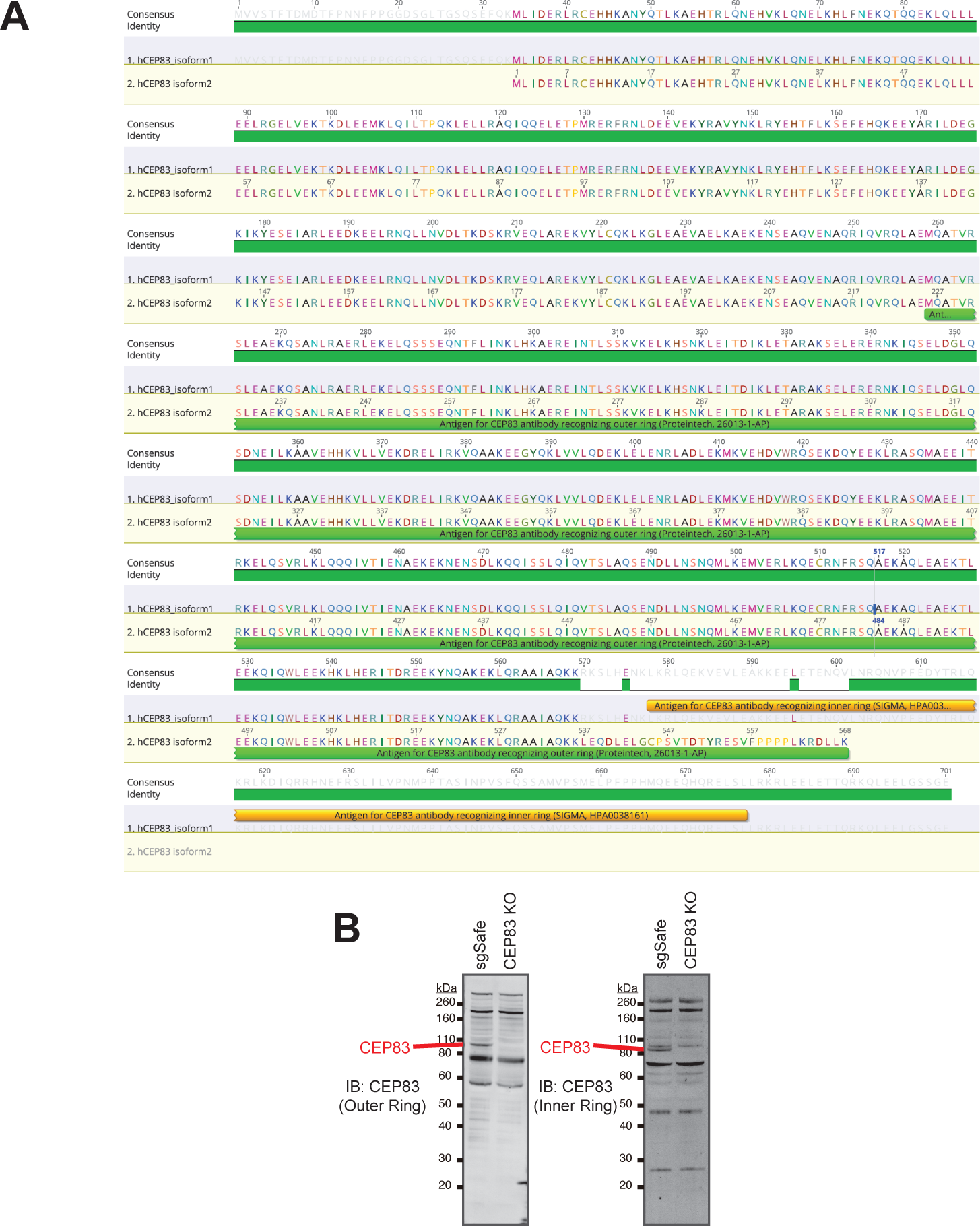
Characterization of the two CEP83 antibodies. A. A DNA sequencing alignment of isoform1 (identifier: Q9Y592-1) and isoform2 (identifier: Q9Y592-2) of human CEP83. The antigen of the antibody (a.a. 226-568 of the isoform 2) that recognizes the outer ring of CEP83 (cat#26013-1-AP, Proteintech) is shown in green. The antigen of the antibody (a.a. 578-677 of the isoform1) that recognizes the inner ring of CEP83 (cat#HPA0038161, SIGMA Aldrich) is shown in orange. B. Immunoblot analysis of CEP83 in sgSafe (control) or CEP83 knockout RPE cells. The PVDF membrane detected with the antibody that recognizes the outer ring of CEP83 (cat#26013-1-AP, Proteintech) or the one that recognizes the inner ring of CEP83 (cat#HPA0038161, SIGMA Aldrich) is shown in left or right respectively. Both antibodies specifically detect CEP83 at similar size (marked by red bar).

**Figure 1-figure supplement 3.**
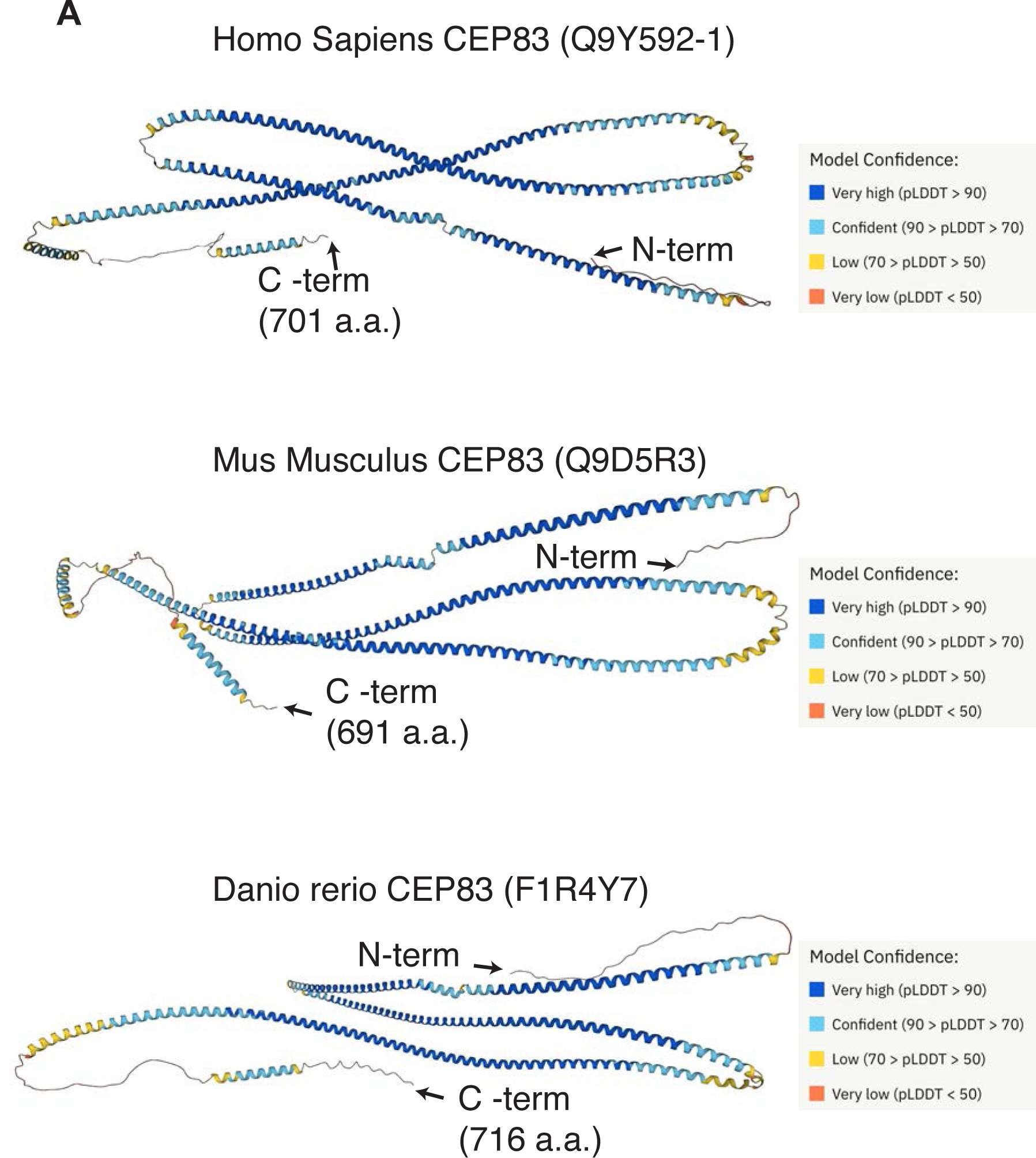
Structural model of CEP83. A. Structural models of CEP83 created by AlphaFold Protein Structure Database. CEP83 structures from the three different species (Homo Sapiens, Mus Musculus, and Danio rerio) show extended alpha helical structures as well as disordered regions at the N- and C-terminus of the protein.

**Figure 1-figure supplement 4.**
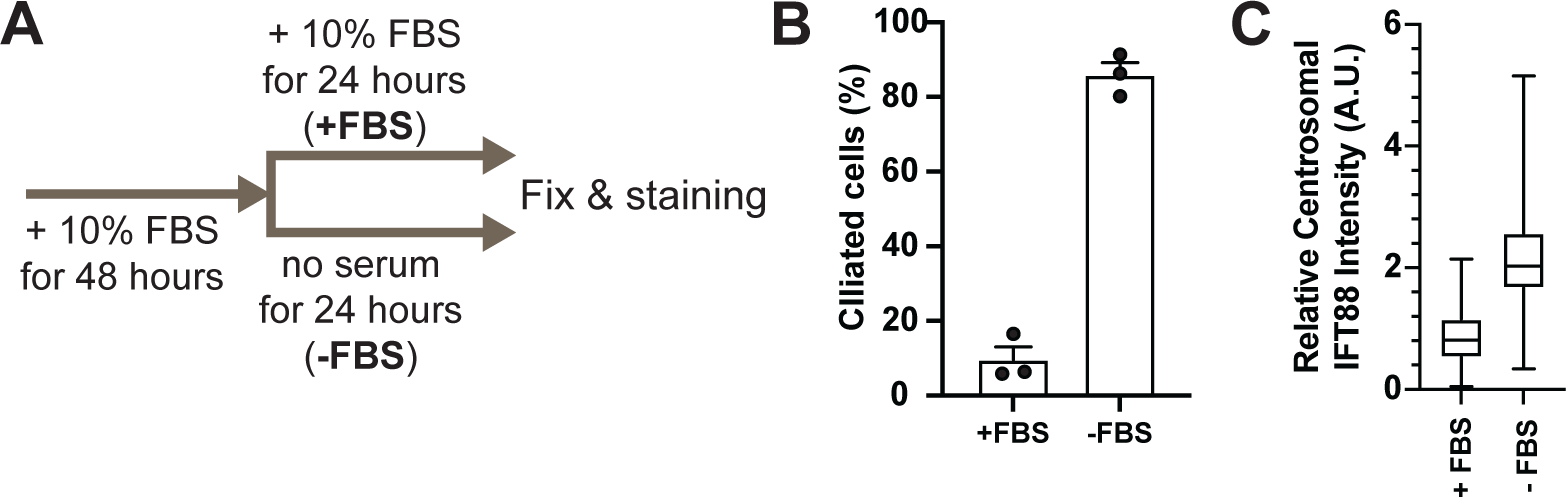
Confirmation of experimental appropriateness of the data shown in Figure 1D. A. The graphical overview of the experimental method used in Figure 1D. B. Quantification of the percentage of the ciliated cells in serum-fed (+FBS) or serum-starved (-FBS) RPE cells. Data obtained from three independent experiments. Each black dot indicates the data from an individual experiment. Error bars represent ± SEM. The raw data, experimental conditions, and detailed statistics are available in Figure 1-figure supplement 4B-Source Data. C. Box plots showing the fluorescent intensity of centrosomal IFT88 in serum-fed (+FBS) or serum-starved (-FBS) RPE cells. The relative fluorescence signal intensity compared with the average of the serum-fed cells is shown. The data is combined from two independent experiments. The raw data and experimental conditions are available in Figure 1-figure supplement 4C-Source Data.

**Figure 2-figure supplement 1.**
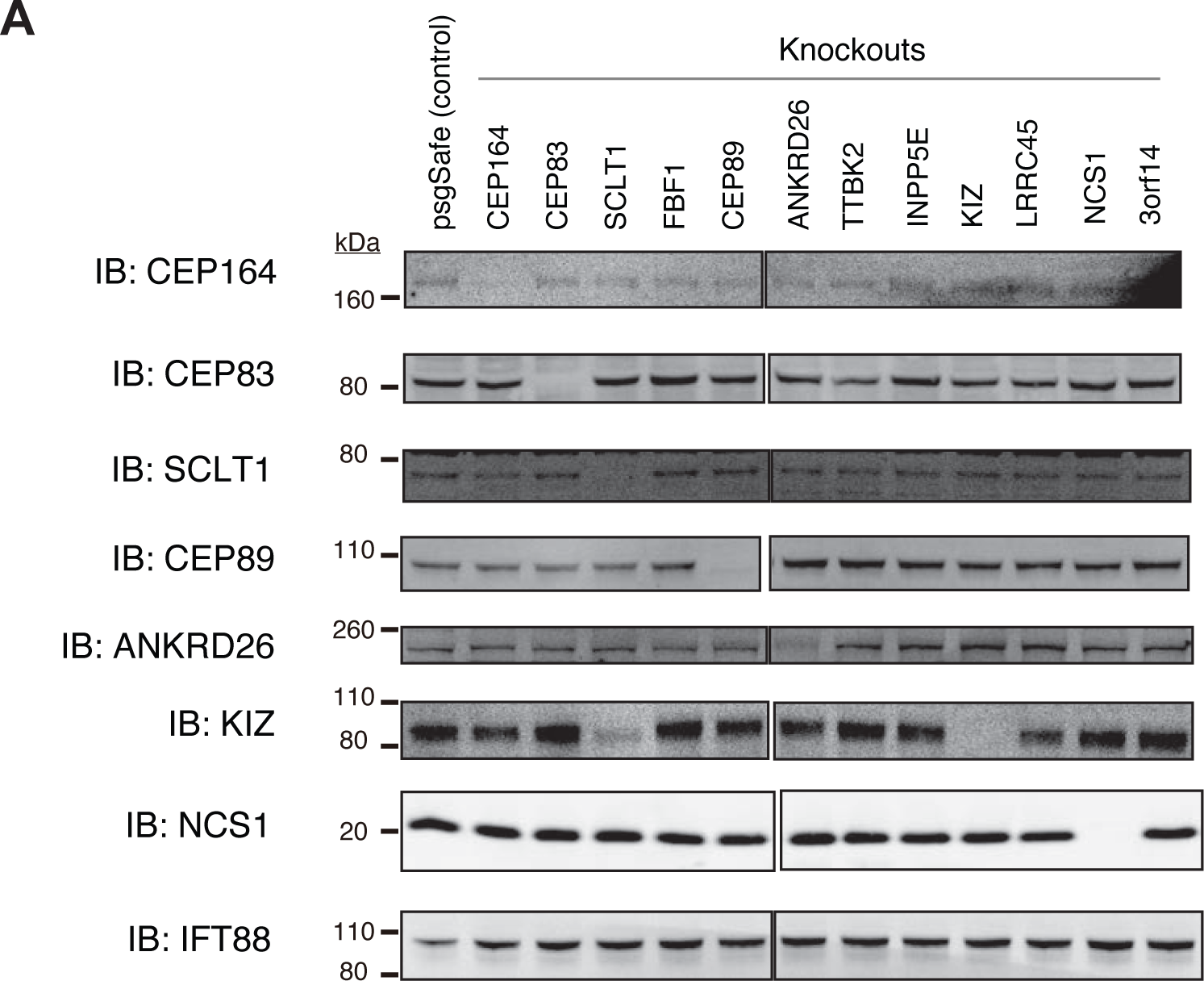
The expression level of distal appendage proteins in the individual distal appendage knockouts. A. Immunoblot (IB) analysis of indicated distal appendage proteins in indicated distal appendage knockout RPE cells grown to confluent (without serum starvation). The expression of distal appendage proteins is generally not affected by other distal appendage proteins except the dramatic reduction of KIZ expression in SCLT1 knockout cells.

**Figure 2-supplementary table 1.**
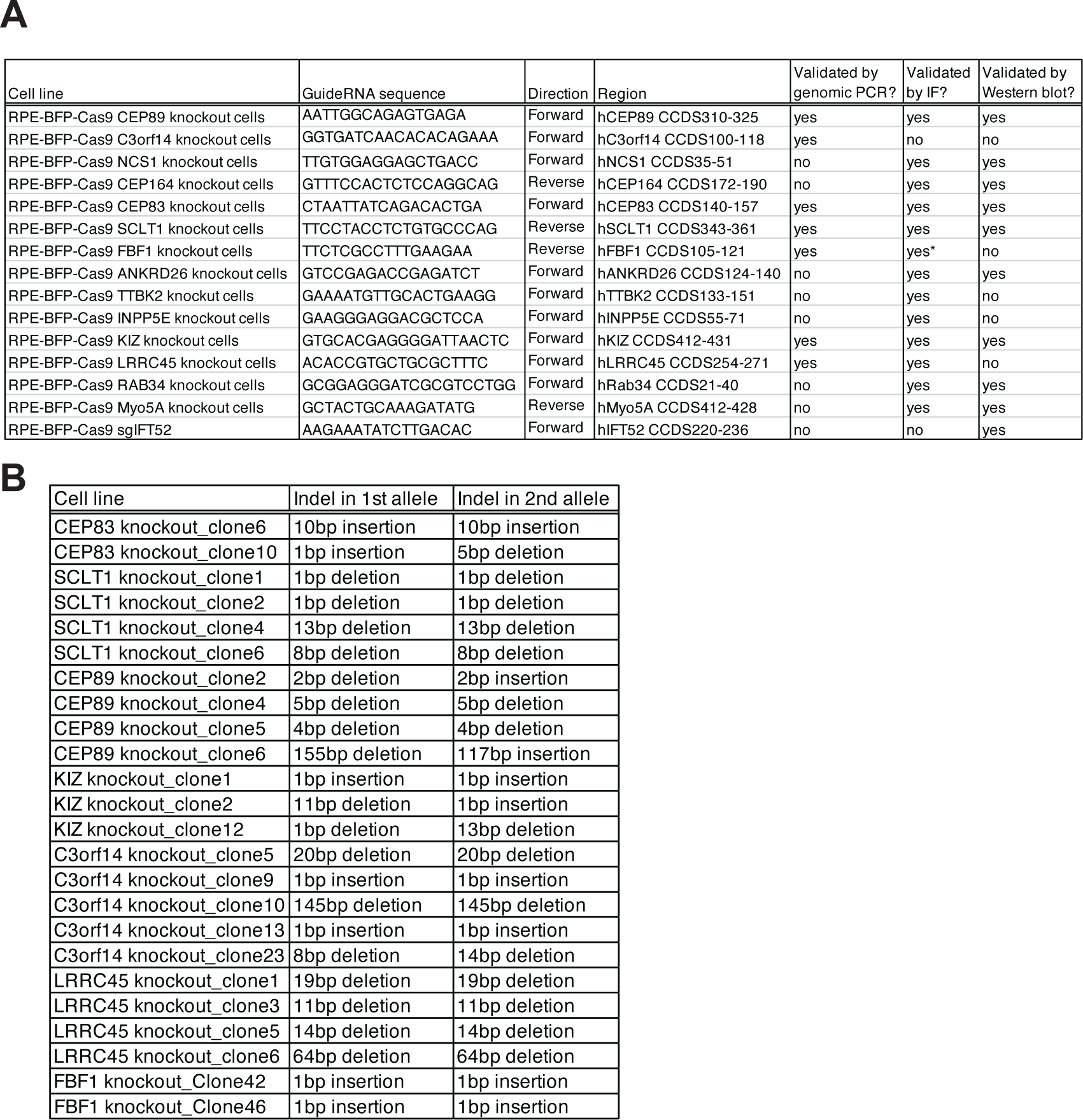
Summary of CRISPR knockout cells used in this paper. A. A table summarizing the target sequence of sgRNAs, direction of the target sequence in the genome, the targeting region within the consensus coding DNA sequence (CCDS), and the methods used for validation of gene depletion. B. A table showing insertion/deletion harbored by each single cell clone of indicated knockout cells. Each insertion/deletion was determined using genomic PCR followed by TIDE analysis.

**Figure 3-figure supplement 1.**
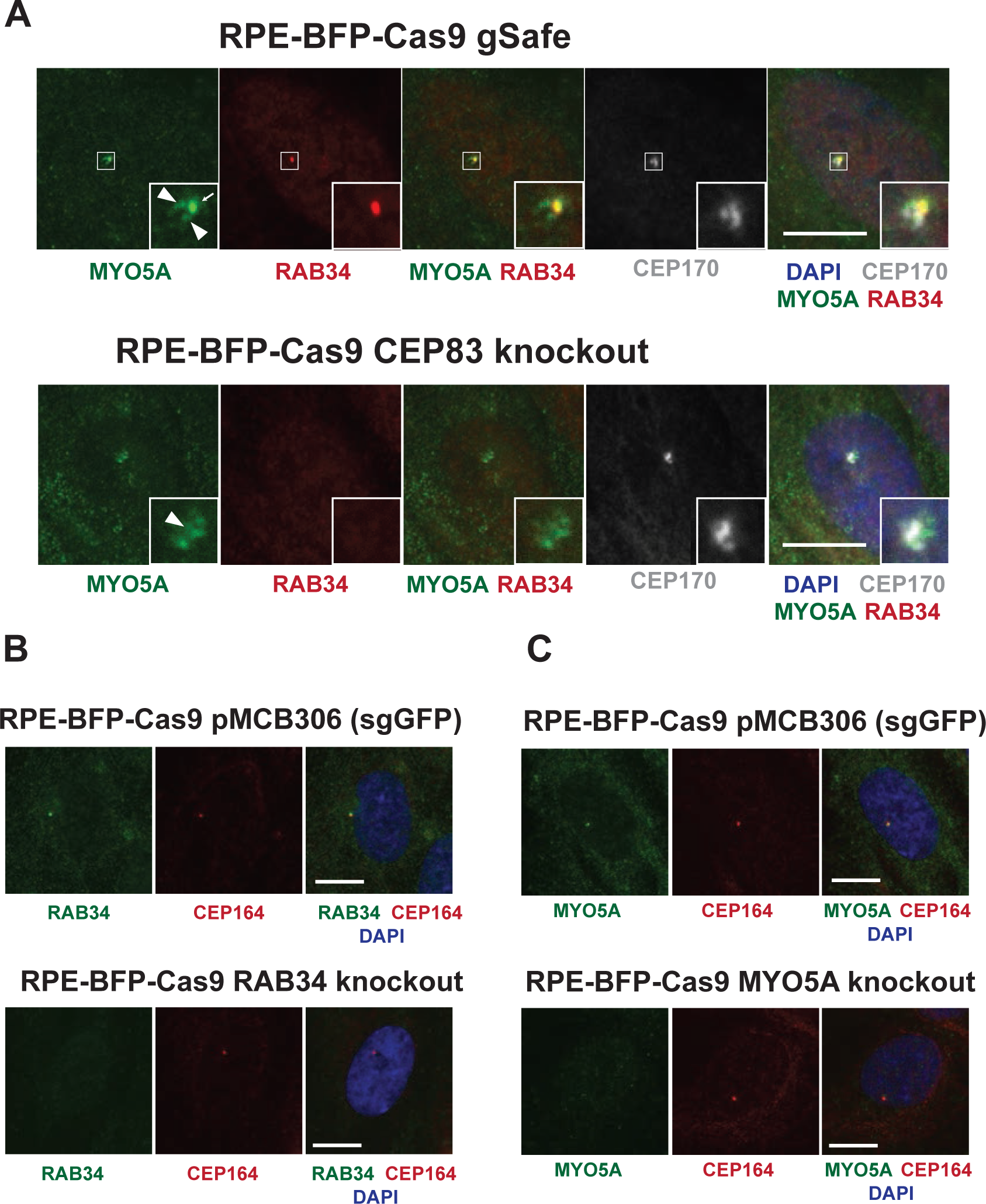
A potential problem of using MYO5A as a ciliary vesicle marker. A. Control (sgSafe) or CEP83 knockout RPE cells were grown to confluent in 10% FBS containing media (serum-fed), fixed, stained with indicated antibodies, and imaged via a wide-field microscopy. Arrow and arrowheads indicate ciliary vesicle and pericentriolar non-ciliary vesicle staining, respectively. Scale bar: 10 *µ*m. Pericentriolar staining observed in MYO5A staining is not evident in RAB34. The ciliary vesicle signal positive for MYO5A and RAB34 is not visible in CEP83 knockout cells, while the pericentriolar staining (arrowhead) persists in the knockout cells. B-C. Control (sgGFP), RAB34 knockout, or MYO5A knockout RPE cells were grown to confluent in 10% FBS containing media (serum-fed), fixed, stained with indicated antibodies, and imaged via a wide-field microscopy. Scale bar: 10 *µ*m. RAB34 and MYO5A signal observed at the mother centrioles (marked by CEP164) and cytoplasm in the control cells are almost completely lost in the respective knockout cells, suggesting that both RAB34 and MYO5A localizes to the mother centrioles and cytoplasm.

**Figure 3-figure supplement 2.**
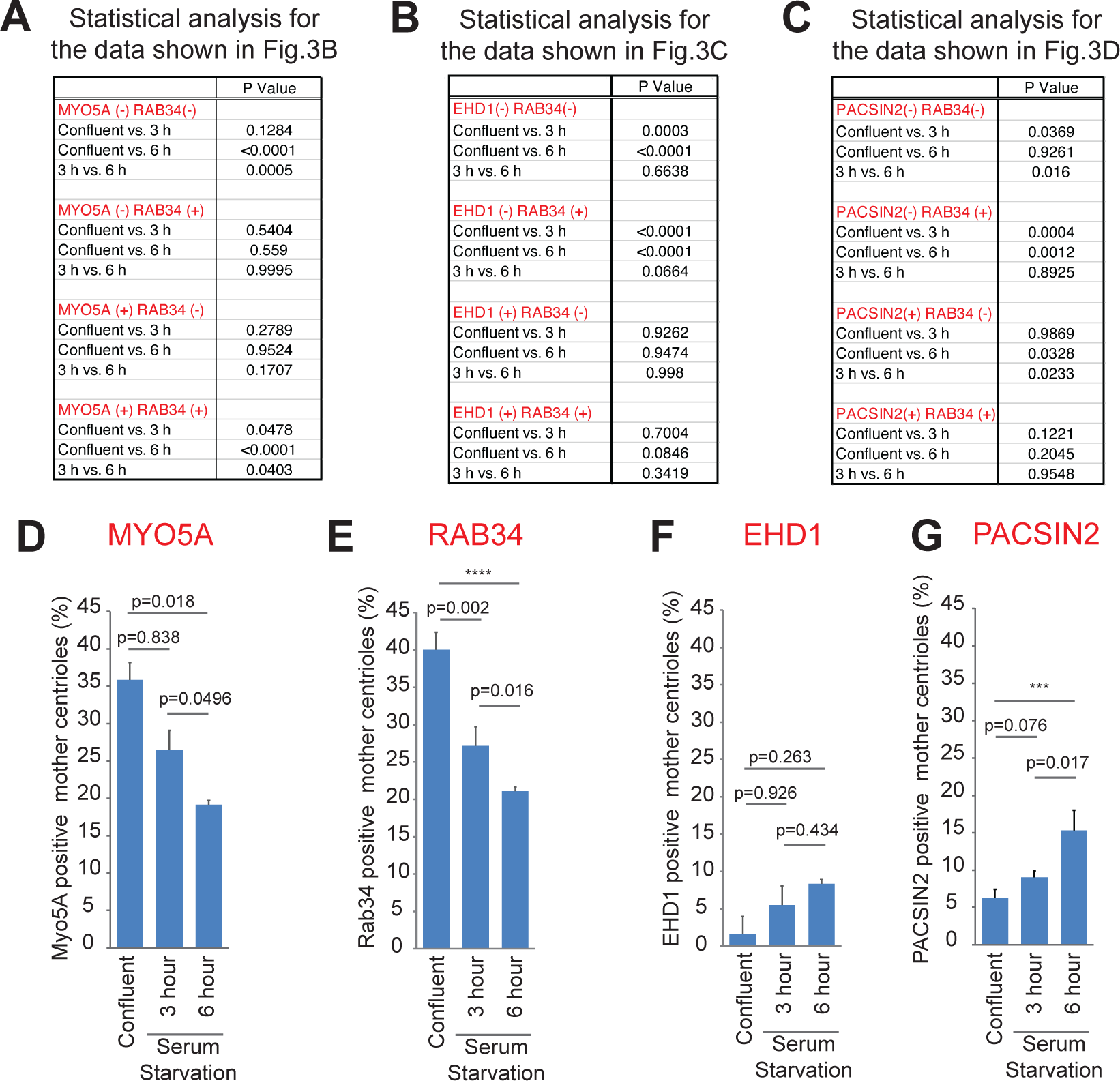
RAB34 and MYO5A are recruited to the mother centriole earlier than EHD1 and PACSIN2. A-C. Key statistics of the data shown in Figure 3B-D. Statistical significance was calculated from two-way ANOVA with Tukey’s multiple comparisons test. Sample numbers and more detailed statistics are available in Figure 3B-Source Data, Figure 3C-Source Data and Figure 3D-Source Data. D-G. Quantification of the percentage of mother centrioles positive for the indicated markers differently processed from the data shown in Figure 3B-D. Statistical significance was calculated from Tukey’s multiple comparisons test. Sample numbers and more detailed statistics are available in Figure 3-figure supplement 2D-Source Data, Figure 3-figure supplement 2F-Source Data, and Figure 3-figure supplement 2G-Source Data.

**Figure 3-figure supplement 3.**
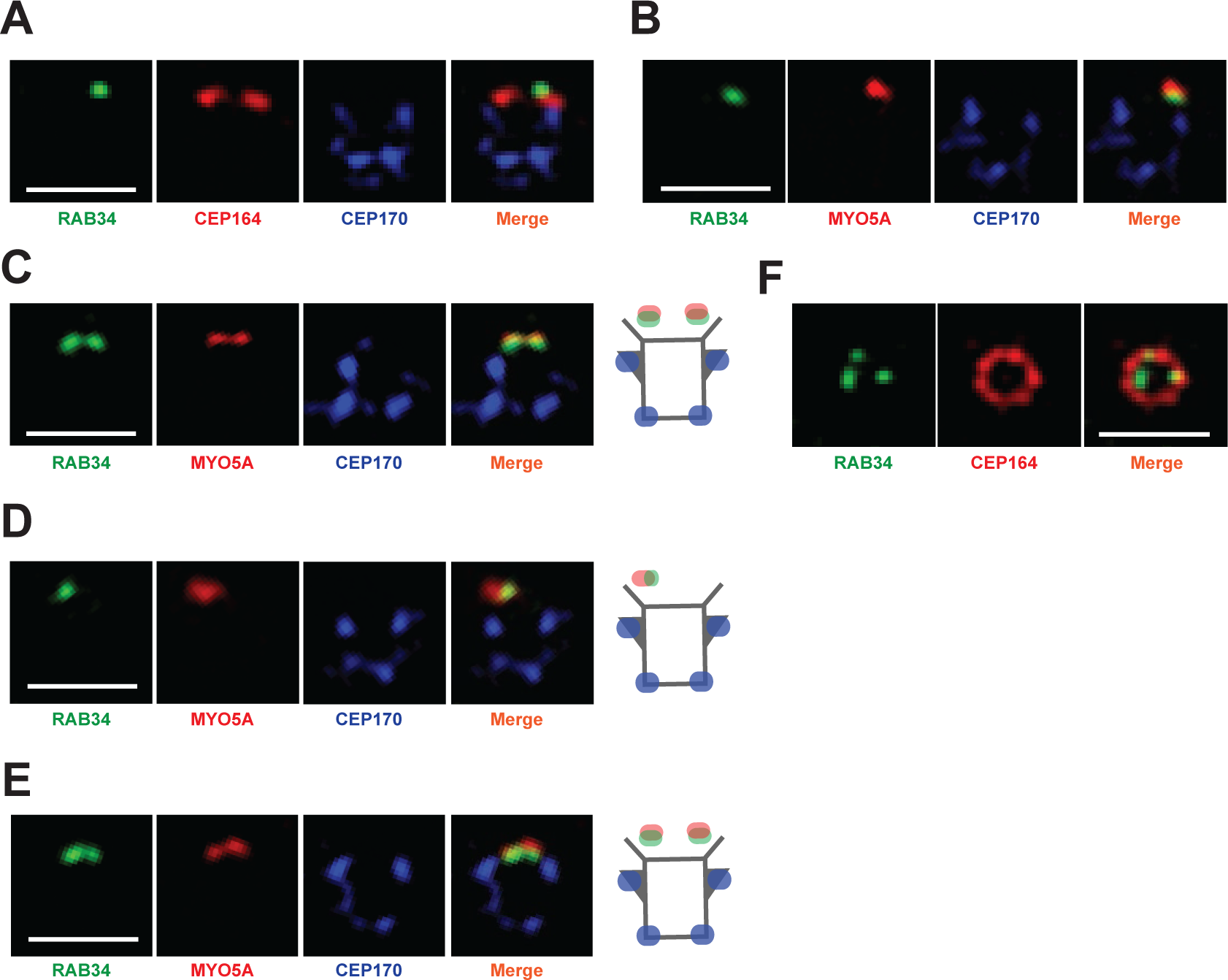
3D structured illumination images of RAB34 and MYO5A. A-B. 3D structured illumination images of indicated distal appendage proteins shown in Figure 3E and G shown in individual channels. Scale bar:1 *µ*m. C-E. Additional 3D structured illumination images of the mother centrioles stained with RAB34 and MYO5A. Scale bar: 1 *µ*m. F. 3D structured illumination image of indicated distal appendage proteins shown in Figure 3F shown in individual channels. Scale bar: 1 *µ*m.

**Figure 3-figure supplement 4.**
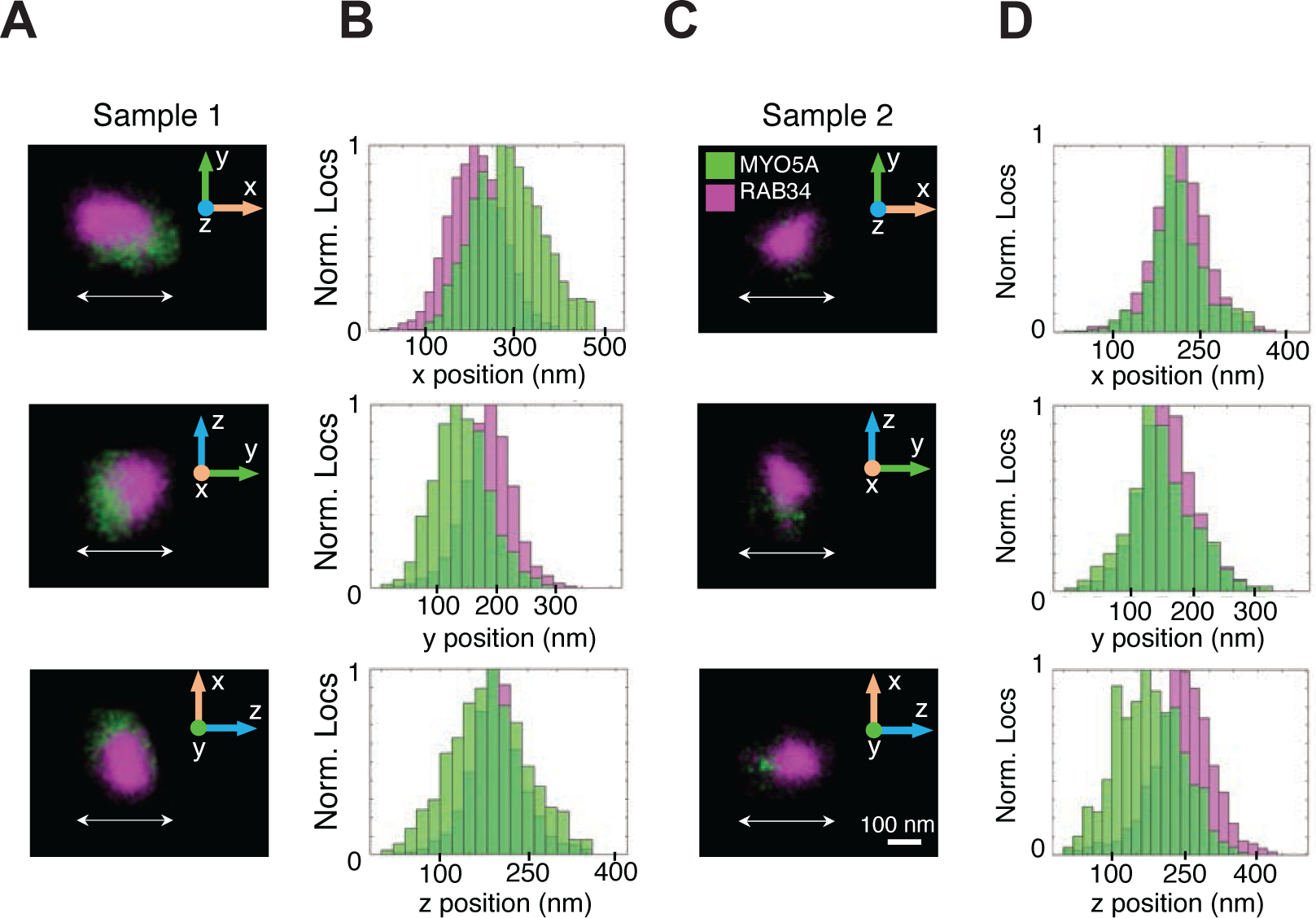
Super-resolution reconstructions of RAB34 and MYO5A manually isolated from the data shown in Figure 3H and 3J with corresponding normalized histograms. A and C. Super-resolution reconstructions of the localizations of MYO5A (green) and RAB34 (magenta). B and D. Normalized histograms of the RAB34 and MYO5A localizations shown in (A) and (C) along each axis with the direction specified by the double-sided arrow shown in (A) and (C).

**Figure 3-figure supplement 5.**
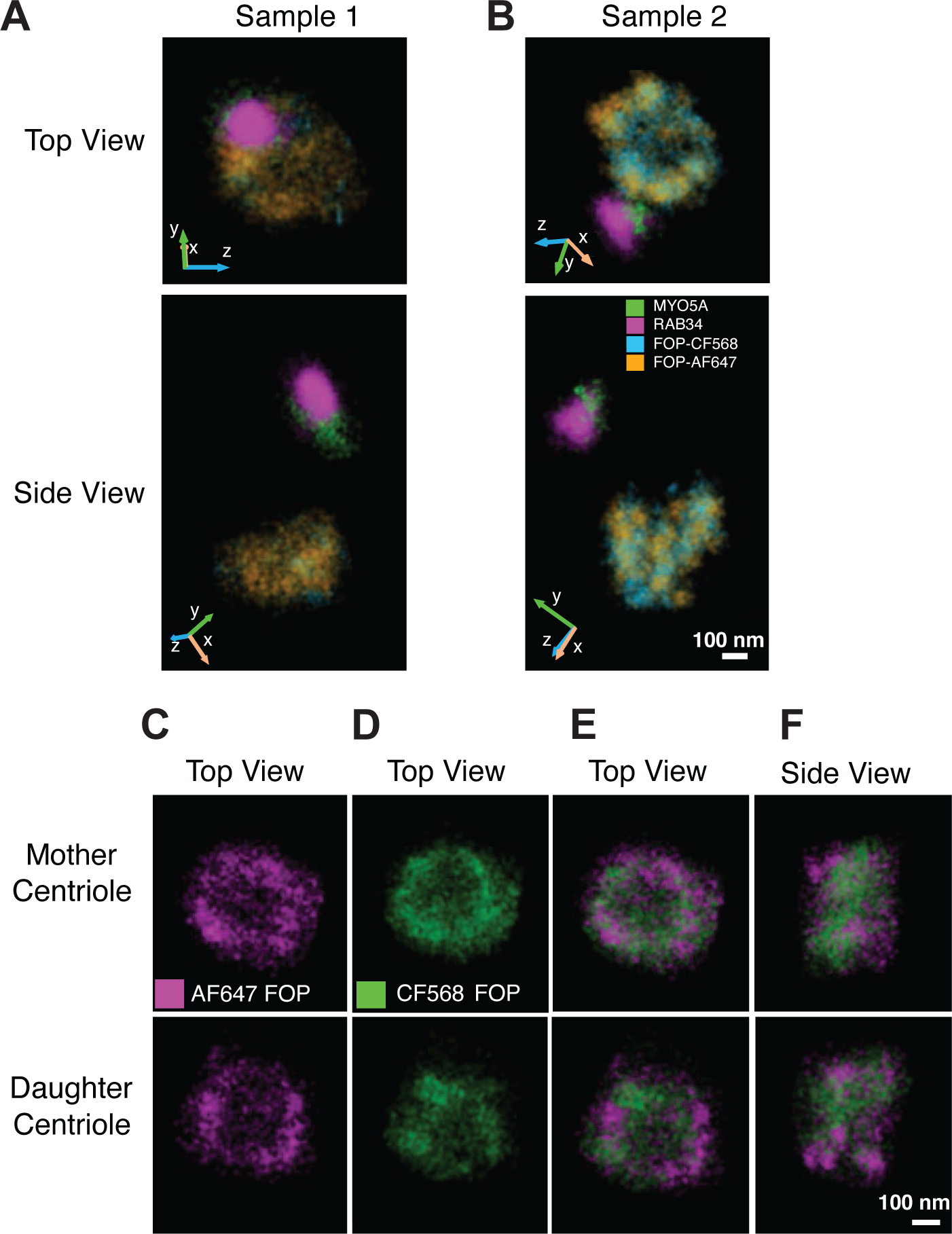
Registration of the 3D single-molecule super-resolution data by imaging of FOP. A and B. 3D single-molecule data from Figure 3H and 3J showing FOP data from the two channels separately. The manually isolated localizations of CF568-labeled FOP (blue) and Alexa Fluor 647 (AF647)-labeled FOP (orange), shown for top and side views, along with CF568-labeled MYO5A (green) and AF647-labeled RAB34 (magenta) for the two samples. Orientations in the microscope 3D space are indicated by the inset axes. The opacities used for visualization in Vutara SRX are as follows: Sample 1, RAB34: 0.03, MYO5A: 0.08, FOP-CF568: 0.05, FOP-AF647: 0.05; sample 2, RAB34: 0.02, MYO5A: 0.09, FOP-CF568: 0.05, FOP-AF647: 0.05. C-F. Example of reconstructions of the FOP ring structures in the two color-channels after channel transformation and cross-correlation of FOP that is labeled in both channels. C-D. Example of the AF647-labeled (magenta) and CF568-labeled (green) FOP structures at the mother and daughter centrioles used for cross-correlation. E-F. The FOP structures from the two channels after channel transformation and data cross-correlation. The opacities for the visualization are set in Vutara SRX to 0.07 for AF647 FOP and 0.02 for CF568 FOP.

**Figure 3-figure supplement 6.**
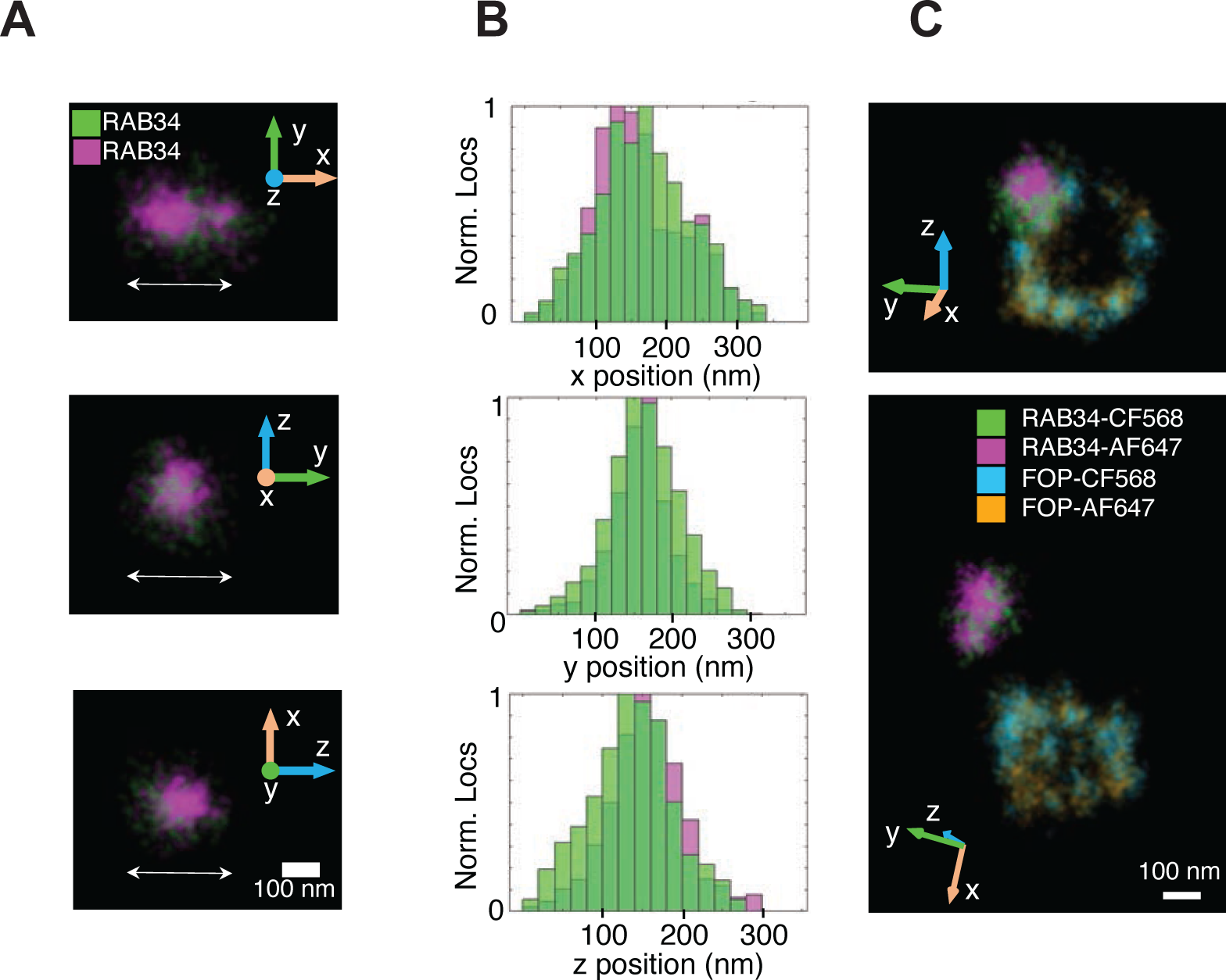
Control of the registration of the 3D single-molecule super-resolution data by imaging of RAB34. A. Super-resolution reconstructions from the localizations of CF568-labeled RAB34 and Alexa Fluor 647 (AF647)-labeled RAB34. The double-sided arrows specify the axes of the histograms shown in (B). B. Offsets between the center of masses of CF568-labeled RAB34 and AF647-labeled RAB34 were found by fitting normalized histograms along each axis to Gaussian functions and calculating the difference between the two peaks. This was done in the x, y, and z directions before finding the total offset in 3D space. The offset between the localizations was found to be 20 nm in this sample, and there were 1705 localizations in the red channel and 1228 localizations in the green channel for RAB34. C. Super-resolution reconstruction of RAB34 and FOP in both channels shown for top view (top) and side view (bottom) after channel registration. Visualizations were rendered in Vutara SRX as 3D Gaussians with 20 nm diameter. The opacities used for visualization are as follows: (A) RAB34-Alexa647: 0.06, RAB34-CF568: 0.06; (C) RAB34-AF647: 0.1, RAB34-CF568: 0.1, FOP-AF647: 0.05, FOP-CF568: 0.05.

**Figure 3-figure supplement 7.**
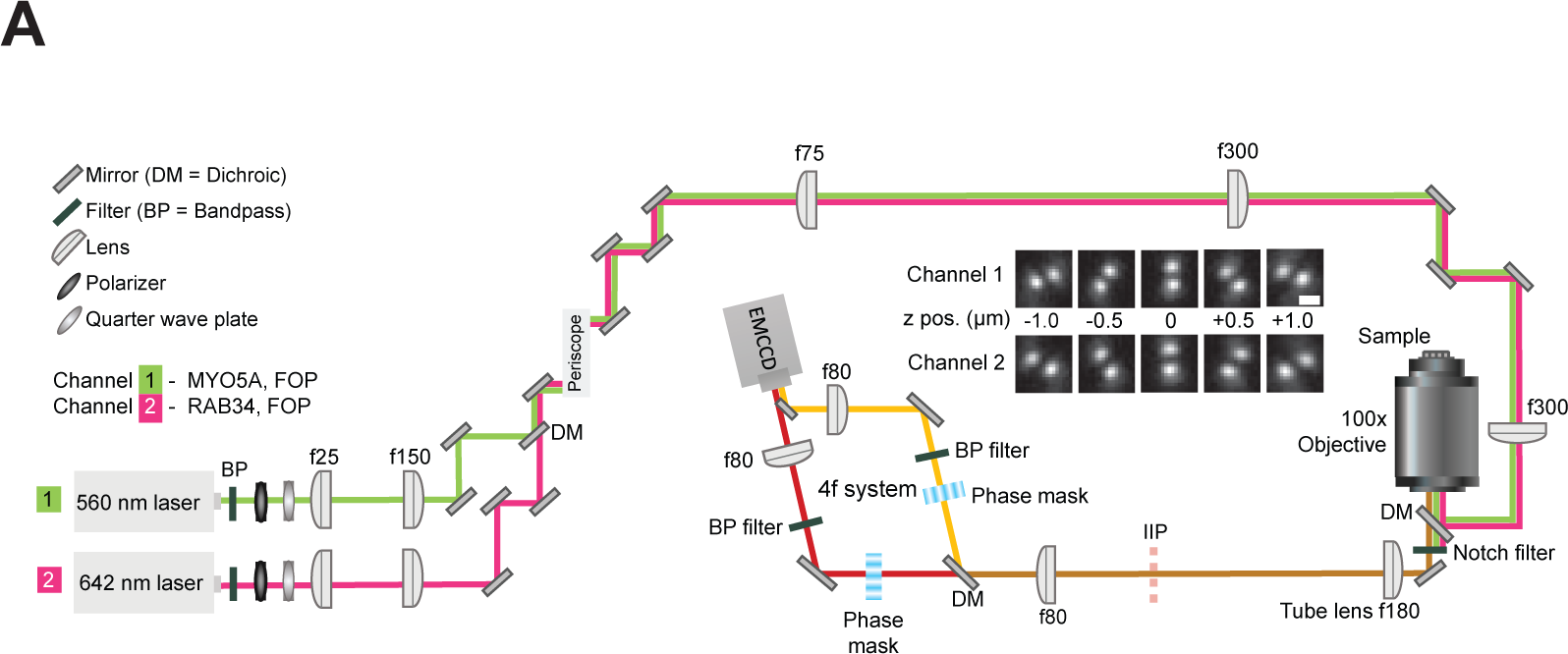
Schematic of the optical setup used to collect 3D single-molecule super-resolution data. Single fluorophores of CF568 and AF647 are excited by 560 nm and 642 nm lasers, respectively, using widefield epi-illumination. A 100x objective lens serves for both illumination and collection of emitted light. A 4f optical relay system is used to image the emitted light, which is split by a dichroic mirror into two emission paths. Transmissive phase masks in the Fourier plane of both paths modulate the emission light, thereby changing the shape of the point spread function (PSF) to that of the double helix PSF, which encodes the 3D position of the emitter. The two paths which are split in the 4f system are imaged onto separate regions of an EMCCD camera. FOP is labeled with both dyes and is imaged in both channels, whereas MYO5A and RAB34 are labeled and imaged in channels 1 and 2, respectively, as indicated above. The inset shows the shape of the double helix PSF in both channels at different axial (z) positions. Scale bar is 1 *µ*m. The schematic is not drawn to scale.

**Figure 4-figure supplement 1.**
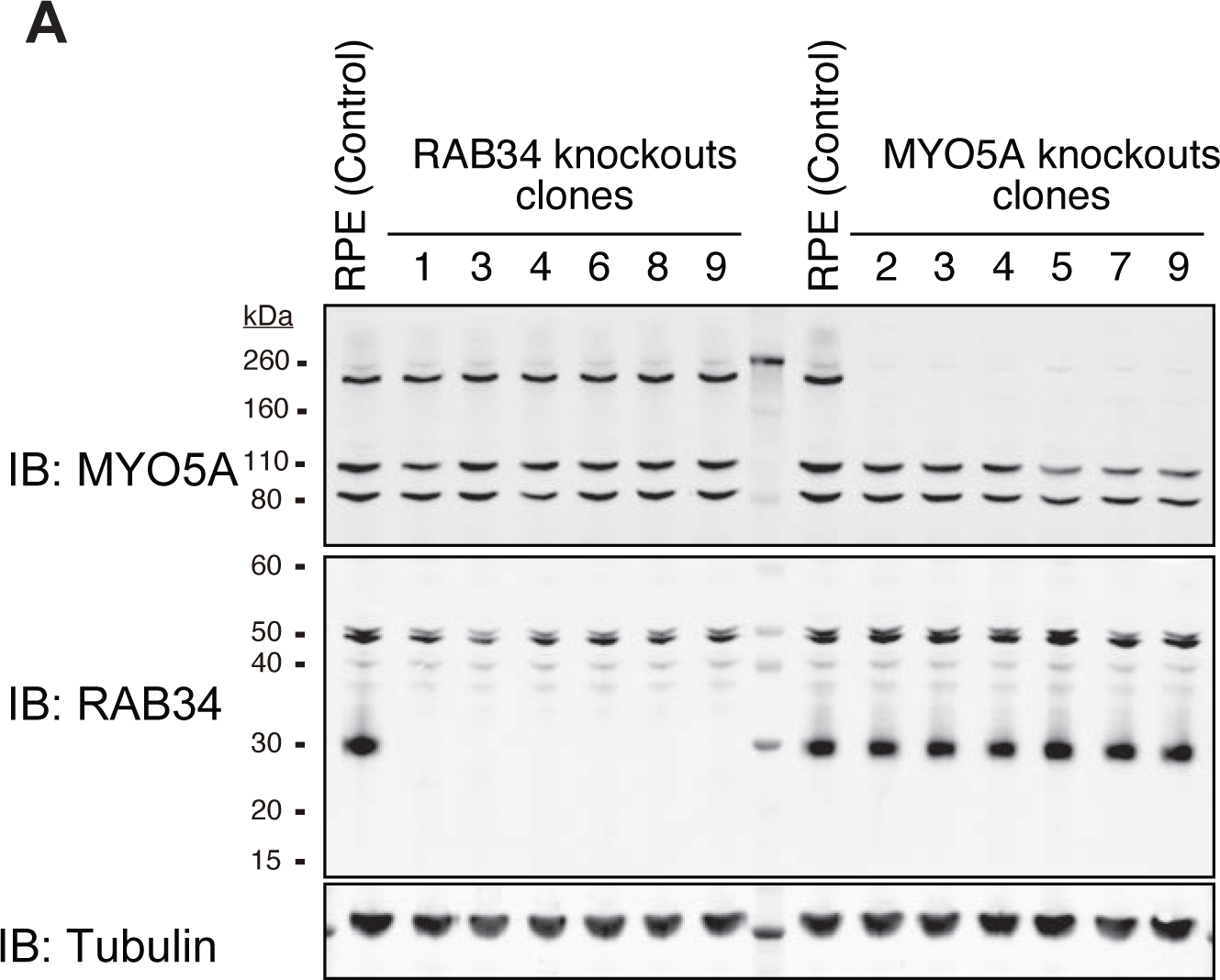
Confirmation of MYO5A and RAB34 knockouts by immunoblot. A. Immunoblot (IB) analysis of MYO5A (IB: MYO5A) and RAB34 (IB: RAB34) in the single cell clones of RAB34 or MYO5A knockout RPE cells. The cells were grown to confluent (without serum starvation), and analyzed by immunoblot. *α*-Tubulin (IB: Tubulin) serves as a loading control.

**Figure 4-figure supplement 2.**
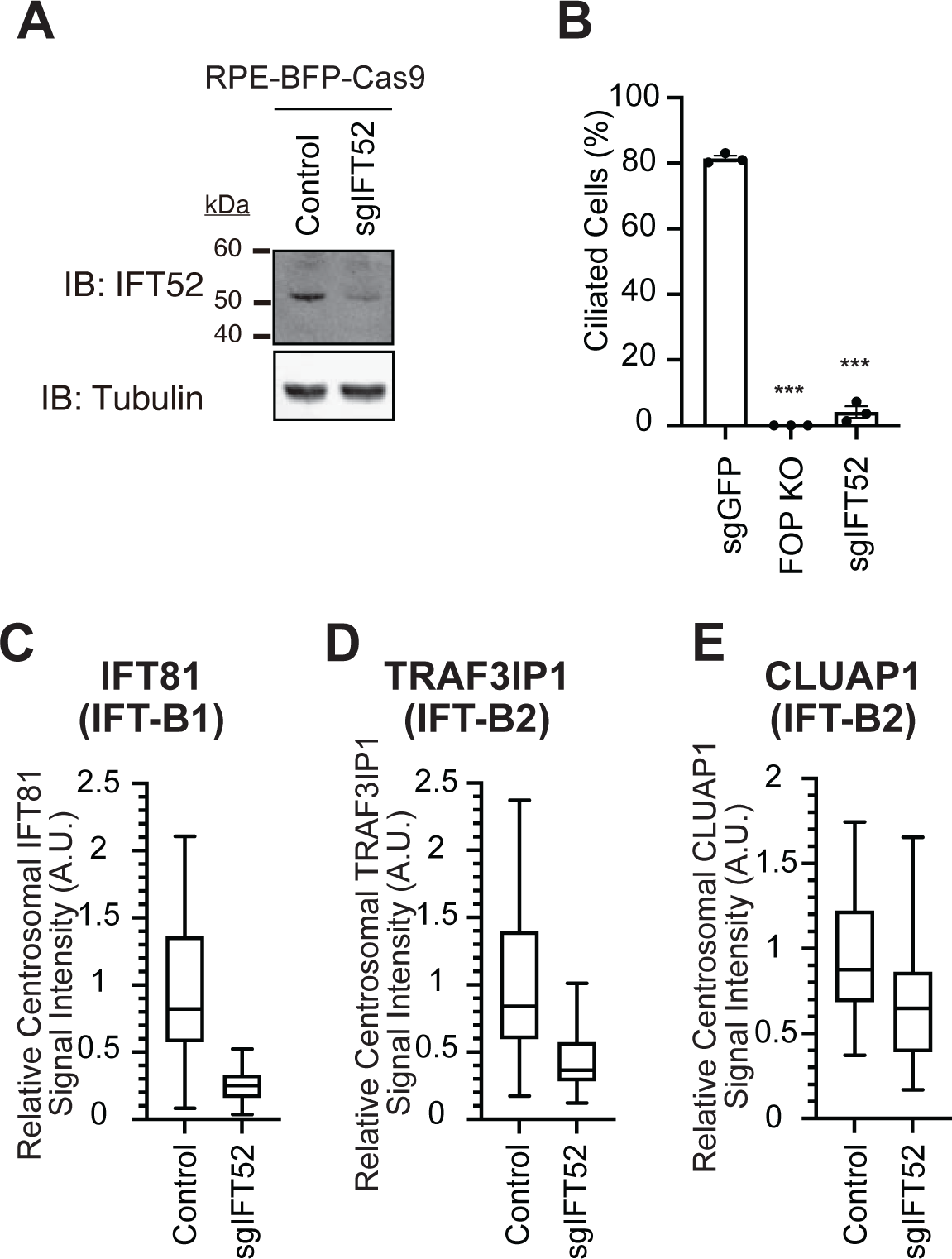
Characterization of IFT52 depleted cell. A. Immunoblot (IB) analysis of IFT52 in either control RPE-BFP-Cas9 or the cells stably expressing sgIFT52. The cells were grown to confluent (without serum starvation), lysed, and analyzed by immunoblot using indicated antibodies. *α*-Tubulin (IB: Tubulin) serves as a loading control. B. Cilium formation assay in control (sgGFP), FOP knockout, or sgIFT52 expressing RPE cells serum starved for 24 hours. Data are averaged from three independent experiments, and each black dot indicates the value from an individual experiment. Error bars represent ± SEM. Statistics obtained through comparing between each knockout and control by Welch’s t-test. The raw data, experimental conditions, and detailed statistics are available in Figure 4-figure supplement 2B-Source Data. C-E. Box plots showing centrosomal signal intensity of IFT81 (C), TRAF3IP1 (D), or CLUAP1 (E) in control (RPE-BFP-Cas9) or RPE cells stably expressing sgIFT52. The relative fluorescence signal intensity compared with the average of the control is shown. At least 40 cells were analyzed per sample. The data from a representative experiment are shown. The raw data and experimental conditions are available in Figure 4-figure supplement 2C-E-Source Data. A.U., arbitrary units; n.s., not significant; *p < 0.05, **p < 0.01, ***p < 0.001

**Figure 5-figure supplement 1.**
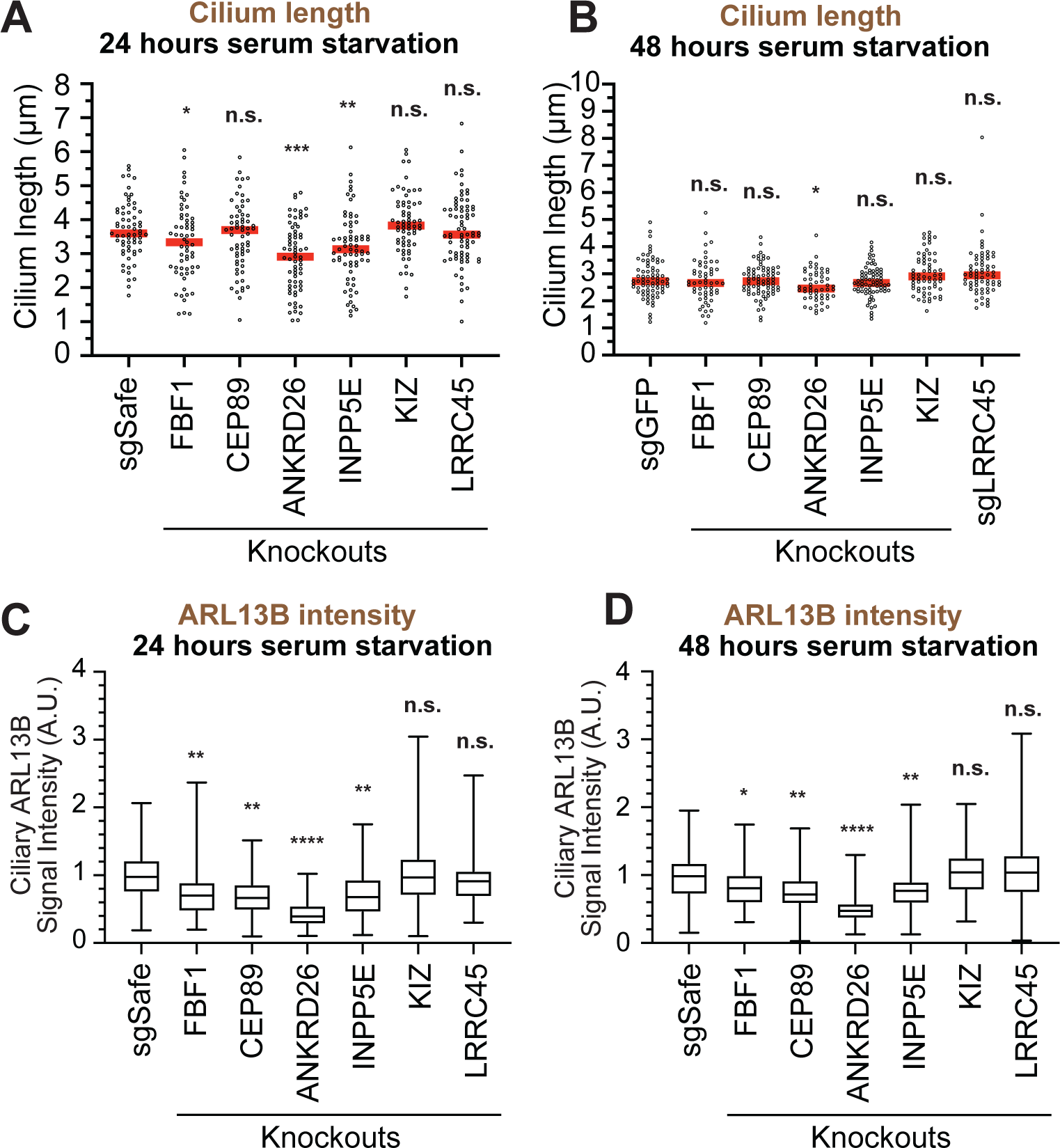
ARL13B intensity was reduced in the various distal appendage knockouts. A-B. Cilium length in control (sgSafe or sgGFP) and indicated knockout RPE cells serum-starved for 24 hours (A) or 48 hours (B). The data from a representative experiment are shown. Each circle indicates the cilium length of the individual cells. Red bars indicate median value. Statistics obtained through comparing between each knockout and control by Welch’s t-test. The raw data, experimental conditions, and detailed statistics are available in Figure 5-figure supplement 2A-B-Source Data. C-D. Box plots showing ciliary signal intensity of ARL13B in control (sgSafe) and indicated knockout RPE cells serum starved for 24 hours (C) or 48 hours (D). The relative fluorescence signal intensity compared with the average of the control is shown. The data from three independent experiments are shown. Statistics obtained through comparing between each knockout and control by nested one-way ANOVA with Dunnett’s multiple comparisons test. The raw data and experimental condition are available in Figure 5-figure supplement 2C-D-Source Data. A.U., arbitrary units; n.s., not significant; *p < 0.05, **p < 0.01, ***p < 0.001

## Source Data

Figure 1A-Source Data. Immunofluorescence conditions in the experiment shown in Figure 1A.

Figure 1C-Source Data. Raw quantification data of the experiment shown in Figure 1C.

Figure 1D-Source Data. Raw quantification data and detailed statistics of the experiment shown in Figure 1D.

Figure 2A-Source Data. Immunofluorescence conditions, and raw quantification data of the experiment shown in Figure 2A.

Figure 2B-Source Data. Immunofluorescence conditions, and raw quantification data of the experiment shown in Figure 2B.

Figure 2C-Source Data. Immunofluorescence conditions, and raw quantification data of the experiment shown in Figure 2C.

Figure 2D-Source Data. Immunofluorescence conditions, and raw quantification data of the experiment shown in Figure 2D.

Figure 2E-Source Data. Immunofluorescence conditions, and raw quantification data of the experiment shown in Figure 2E.

Figure 2F-Source Data. Immunofluorescence conditions, and raw quantification data of the experiment shown in Figure 2F.

Figure 2G-Source Data. Immunofluorescence conditions, and raw quantification data of the experiment shown in Figure 2G.

Figure 2H-Source Data. Immunofluorescence conditions, and raw quantification data of the experiment shown in Figure 2H.

Figure 2I-Source Data. Immunofluorescence conditions, and raw quantification data of the experiment shown in Figure 2I.

Figure 2J-Source Data. Immunofluorescence conditions, and raw quantification data of the experiment shown in Figure 2J.

Figure 2K-Source Data. Immunofluorescence conditions, and raw quantification data of the experiment shown in Figure 2K.

Figure 2L-Source Data. Immunofluorescence conditions, and raw quantification data of the experiment shown in Figure 2L.

Figure 3A, E, F, G-Source Data. Immunofluorescence conditions in the experiment shown in Figure 3A, E, F, and G.

Figure 3B-Source Data. Raw quantification data, immunofluorescence conditions and detailed statistics of the experiment shown in Figure 3B.

Figure 3C-Source Data. Raw quantification data, immunofluorescence conditions and detailed statistics of the experiment shown in Figure 3C.

Figure 3D-Source Data. Raw quantification data, immunofluorescence conditions and detailed statistics of the experiment shown in Figure 3D.

Figure 4A-Source Data. Raw quantification data, immunofluorescence conditions and detailed statistics of the experiment shown in Figure 4A.

Figure 4C-Source Data. Raw quantification data and detailed statistics of the experiment shown in Figure 4C.

Figure 4D-Source Data. Raw quantification data, immunofluorescence conditions and detailed statistics of the experiment shown in Figure 4D.

Figure 4E-Source Data. Raw quantification data and immunofluorescence conditions of the experiment shown in Figure 4E.

Figure 4F-Source Data Raw quantification data and immunofluorescence conditions of the experiment shown in Figure 4F.

Figure 4G-Source Data. Raw quantification data and immunofluorescence conditions of the experiment shown in Figure 4G.

Figure 4H-Source Data. Raw quantification data and immunofluorescence conditions of the experiment shown in Figure 4H.

Figure 4I-Source Data. Raw quantification data and immunofluorescence conditions of the experiment shown in Figure 4I.

Figure 4J-Source Data. Raw quantification data and immunofluorescence conditions of the experiment shown in Figure 4J.

Figure 4K-Source Data. Raw quantification data, immunofluorescence conditions and detailed statistics of the experiment shown in Figure 4K.

Figure 5A-Source Data. Raw quantification data, immunofluorescence conditions and detailed statistics of the experiment shown in Figure 5A.

Figure 5B-Source Data. Raw quantification data, immunofluorescence conditions and detailed statistics of the experiment shown in Figure 5B.

Figure 5C-Source Data. Raw quantification data, immunofluorescence conditions and detailed statistics of the experiment shown in Figure 5C.

Figure 5D-Source Data. Raw quantification data, immunofluorescence conditions and detailed statistics of the experiment shown in Figure 5C.

Figure 5E-Source Data. Raw quantification data and immunofluorescence conditions of the experiment shown in Figure 5E.

Figure 5F-Source Data. Raw quantification data and immunofluorescence conditions of the experiment shown in Figure 5F.

Figure 1-figure supplement 2B-Source Data. The original files of the full raw unedited blots shown in Figure 1-figure supplement 2B.

Figure 1-Figure Supplement 4B-Source Data. Raw quantification data, immunofluorescence conditions and detailed statistics of the experiment shown in Figure 1-Figure supplement 4B.

Figure 1-Figure Supplement 4C-Source Data. Raw quantification data and immunofluorescence conditions of the experiment shown in Figure 1-Figure supplement 4C.

Figure 2-figure supplement 1A-Source Data. The original files of the full raw unedited blots shown in Figure 2-figure supplement 1A.

Figure 3-Figure Supplement 1-Source Data. Raw quantification data and immunofluorescence conditions of the experiment shown in Figure 3-Figure supplement 1A, B and C.

Figure 3-Figure Supplement 2D and E-Source Data. Raw quantification data and detailed statistics of the experiment shown in Figure 3-Figure supplement 2D and E.

Figure 3-Figure Supplement 2F-Source Data. Raw quantification data and detailed statistics of the experiment shown in Figure 3-Figure supplement 2F.

Figure 3-Figure Supplement 2G-Source Data. Raw quantification data and detailed statistics of the experiment shown in Figure 3-Figure supplement 2G.

Figure 3-Figure Supplement 3-Source Data. Immunofluorescence conditions in the experiment shown in Figure 3-Figure Supplement 3A-F.

Figure 4-figure supplement 1A-Source Data. The original files of the full raw unedited blots shown in Figure 4-figure supplement 1A.

Figure 4-figure supplement 2A-Source Data. The original files of the full raw unedited blots shown in Figure 4-figure supplement 2A.

Figure 4-Figure Supplement 2B-Source Data. Raw quantification data, immunofluorescence conditions and detailed statistics of the experiment shown in Figure 4-Figure supplement 2B.

Figure 4-Figure Supplement 2C-Source Data. Raw quantification data and immunofluorescence conditions of the experiment shown in Figure 4-Figure supplement 2C.

Figure 4-Figure Supplement 2D-Source Data. Raw quantification data and immunofluorescence conditions of the experiment shown in Figure 4-Figure supplement 2D.

Figure 4-Figure Supplement 2E-Source Data. Raw quantification data and immunofluorescence conditions of the experiment shown in Figure 4-Figure supplement 2E.

Figure 5-Figure Supplement 1A-Source Data. Raw quantification data, immunofluorescence conditions and detailed statistics of the experiment shown in Figure 5-Figure supplement 1A.

Figure 5-Figure Supplement 1B-Source Data. Raw quantification data, immunofluorescence conditions and detailed statistics of the experiment shown in Figure 5-Figure supplement 1B.

Figure 5-Figure Supplement 1C-Source Data. Raw quantification data, immunofluorescence conditions and detailed statistics of the experiment shown in Figure 5-Figure supplement 1C.

Figure 5-Figure Supplement 1D-Source Data. Raw quantification data, immunofluorescence conditions and detailed statistics of the experiment shown in Figure 5-Figure supplement 1D.

Source Data-the original files of the full raw unedited immunoblot with label.

Source Data-Macro for measuring fluorescent intensity of centrosomal proteins.

Source Data-Macro for measuring fluorescent intensity of ciliary proteins.

Source Data-List of cell lines used in this paper.

Source Data-List of the antibodies -Distal appendage network-.

Source Data-Primers used for genomic PCR

