## Supplementary material for "A hierarchical pathway for assembly of the distal appendages that organize primary cilia": Source Data- the original files of the full raw unedited immunoblot

Figure 1-figure supplement 2B_CEP83

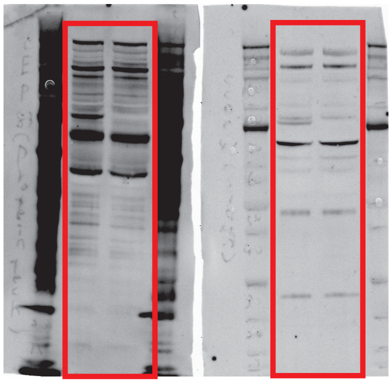

Outer Ring (Proteintech)

Inner Ring (SIGMA)

Figure 2-figure supplement 1A_CEP164

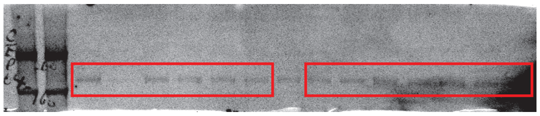

Figure 2-figure supplement 1A_CEP89

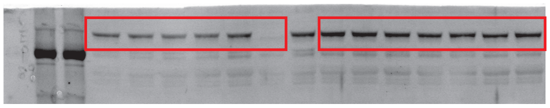

Figure 2-figure supplement 1A_CEP83

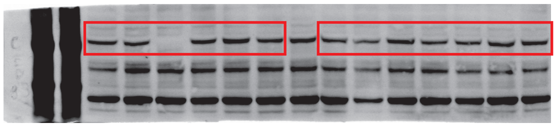

Figure 2-figure supplement 1A_ANKRD26

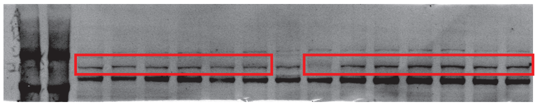

Figure 2-figure supplement 1A_KIZ

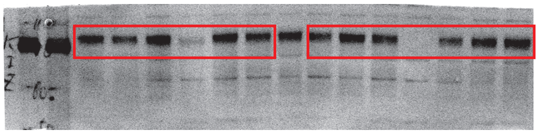

Figure 2-figure supplement 1A_NCS1

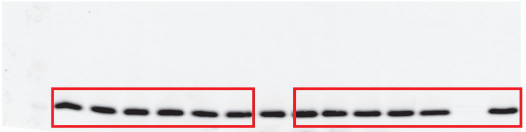

Figure 2-figure supplement 1A_SCLT1

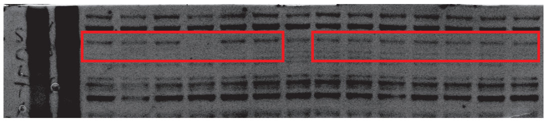

Figure 2-figure supplement 1A_IFT88

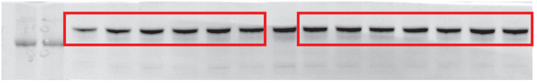

Figure 4-figure supplement 1A_RAB34

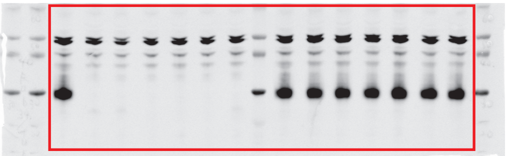

Figure 4-figure supplement 1A_MYO5A

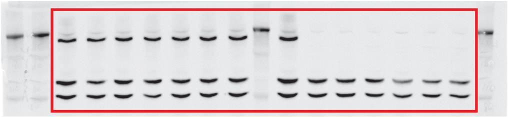

Figure 4-figure supplement 1A_Tubulin

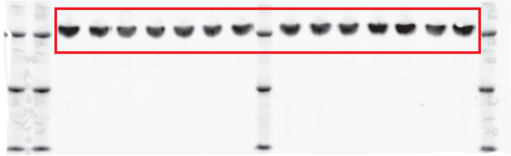

Figure 4-figure supplement 2A_IFT52

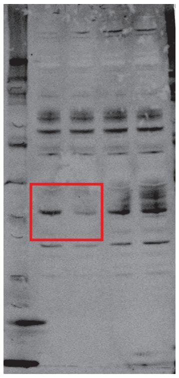

Figure 4-figure supplement 2A_Tubulin

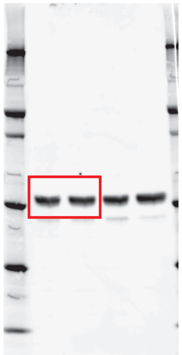

Kanie et al., Resource-full blot
